# Piccolo Regulates Secretion of the Extracellular Matrix Components Brevican and Tenascin R from Astrocytes to Drive Synapse Formation: Implications for Pontocerebellar Hypoplasia Type 3 (PCH3)

**DOI:** 10.1101/2025.07.03.662734

**Authors:** Maryam Mohamaddokht, Joanne Falck, Sanra Öktem, Patrizia Rizzu, Katharina Friedrich, Peter Heutink, Christian Rosenmund, Craig Garner, Dietmar Schmitz, Frauke Ackermann

**Author notes:** Correspondence to Frauke Ackermann.

## Abstract

**Background:** Astrocytes are crucial for CNS health, for instance via the secretion of extracellular matrix (ECM) components that are vital for synapse formation and maturation. While the scaffolding protein Piccolo is known for its role at synapses, its function in astrocytes and contribution to neurodegenerative disorders like Pontocerebellar Hypoplasia Type 3 (PCH3) are largely unknown. Understanding these mechanisms is key to elucidating PCH3 pathology.

**Methods:** We used a multi-faceted approach with a *Pclo^gt/gt^*rat model. Methods included RNA-sequencing for gene expression and GO analysis. Immunohistochemistry and immunocytochemistry assessed Piccolo localization, ECM component (Brevican [Bcan], Tenascin-R [TNR]) levels, distribution, and secretion in brain sections and primary astrocytes. Golgi morphology was evaluated via GM130 staining. Neuronal network formation and function were investigated by co-culturing wild-type neurons with *Pclo^wt/wt^* or *Pclo^gt/gt^* astrocytes in a Banker setup, assessing synapse density (immunostaining, RRP electrophysiology) and spontaneous activity (mEPSCs, mIPSCs). Astrocyte-conditioned media (ACM) experiments determined secreted factor roles.

**Results:** RNA-seq showed significant DEG increases in older versus young *Pclo^gt/gt^* rats (P25 vs. P5), which were strongly enriched in cell communication, signaling, and ECM GO terms. We identified a novel astrocyte-specific Piccolo isoform that partially localizes at the Golgi. Piccolo gene trap transposon mutation led to impaired ECM secretion (reduced extracellular Bcan, altered TNR) from astrocytes, correlating with a fragmented Golgi in *Pclo^gt/gt^* astrocytes. Functionally, *Pclo^gt/gt^* astrocytes significantly reduced synapse density and altered intrinsic activity (increased mEPSC frequency) in neuronal networks. This synaptic deficit was substantially rescued by *Pclo^wt/wt^*astrocyte conditioned media (ACM).

**Conclusion:** Our findings reveal a critical, unrecognized role for Piccolo in regulating astrocytic ECM secretion, essential for proper neuronal network formation and activity. Astrocytic Piccolo dysfunction disrupts this process, causing impaired synaptogenesis and altered intrinsic network activity, providing a novel cellular and molecular mechanism for neurodegenerative diseases like PCH3 and highlights astrocytes as potential therapeutic targets.

## Background

The development, function and health of the brain rely on the intricate interplay between various cell types; in addition to oligodendrocytes and microglia, neurons and astrocytes play essential roles. Neurons serve as the primary signaling units, transmitting electrical and chemical messages, whereas astrocytes provide crucial support. One of the key responsibilities of astrocytes is the regulation of the brain ECM. They secrete ECM components such as glycoproteins, hyaluronan and proteoglycans such as brevican (BCAN), neurocan (NCAN) and tenascin-R (TNR) (Wiese, Karus, and Faissner 2012), as well as synaptogenic factors, which together are necessary for proper neuronal network formation and synaptogenesis (Faissner et al. 2010). In addition to ECM regulation, astrocytes support synaptic transmission by clearing neurotransmitters and releasing gliotransmitters, maintaining extracellular homeostasis by regulating ion concentrations, ensuring blood‒brain barrier integrity, promoting neurogenesis, and providing metabolic support by supplying neurons with essential nutrients (Hart and Karimi-Abdolrezaee 2021; Verkhratsky et al. 2021).

The coordinated interactions between neurons and astrocytes ensure proper neural development, synaptic plasticity, and overall brain function. Neuronal and astrocyte malfunction are the main underlying causes of neurodegenerative diseases. For example, astrocyte dysfunction and ECM abnormalities have been increasingly linked to neurodegenerative diseases, where reactive astrocytes contribute to inflammation, oxidative stress, and neuronal damage. In Alzheimer’s disease (AD), astrocytes promote neuroinflammation and amyloid-beta plaque accumulation (Henstridge, Hyman, and Spires-Jones 2019)), whereas in Parkinson’s disease (PD), their impaired support leads to oxidative stress and dopaminergic neuron degeneration (Booth, Hirst, and Wade-Martins 2017).

The severe early-onset neurodegenerative disorder Pontocerebellar Hypoplasia Type 3 (PCH3) is characterized by profound cerebellar and brainstem hypoplasia. Affected children exhibit developmental delay, intellectual disability, defects in motor control, progressive microcephaly with brachycephaly, seizures, hypotonia with hyperreflexia, and short stature (Rajab et al. 2003). Intriguingly, whole-genome studies linked a single-nucleotide polymorphism (SNP) within the *PCLO (Piccolo)* gene with PCH3 (Ahmed et al. 2015). Piccolo is a presynaptic scaffolding protein that is best known for its role in actin dynamics, synaptic vesicle (SV) release/recycling, maintenance of synapse integrity, and regulation of protein degradation (Wagh et al. 2015; Terry-Lorenzo et al. 2016; Leal-Ortiz et al. 2008; Waites et al. 2011, 2013; Ackermann et al. 2019; Steven D. Fenster and Garner 2002). However, Piccolo is present not only within presynaptic terminals but also in axonal growth cones (Steven D. Fenster et al. 2003) and nonneuronal cells, such as pancreatic islet cells, where it plays a role in insulin secretion (Fujimoto et al. 2002).

A recently characterized PCH3 animal model, harboring a Piccolo gene trap mutation (*Pclo^gt/gt^*), revealed core phenotypes of the disease. These include a thinner cerebral cortex and reduced cell numbers in the cerebellum and pons. Behavioral and functional studies have demonstrated that *Pclo^gt/gt^* rats exhibit impaired motor control (Falck et al. 2020) as well as seizures (A. Medrano et al. 2018), a hallmark of the disease. Furthermore, subcellular analysis of *Pclo^gt/gt^* rats revealed defects in the formation of synapses, as the morphology of mossy fiber terminals (MFs) is dramatically altered. MFs are smaller and disorganized in addition the GABA_A_ɑ6 subunit is downregulated, which is predicted to cause changes in cerebellar network activity, possibly accounting for the motor deficits observed in *Pclo^gt/gt^* rats and patients with PCH3 (Falck et al. 2020). Although this initial study provides hints regarding potential mechanisms underlying PCH3, how Piccolo genetrap mutation contributes to disorganized synapses and contributes to disease pathogenesis remains unclear.

In this study, we identified a novel role for Piccolo in astrocytic ECM secretion and its impact on neuronal network formation. Our findings demonstrate that Piccolo is present in brain astrocytes, where it regulates the secretion of the ECM components BCAN and TNR. Subsequently, impaired BCAN/TNR secretion in a *Pclo^gt/gt^* astrocyte - neuron coculture model led to reduced synapse formation and increased spontaneous activity. These data provide new insights into how Piccolo malfunction contributes to PCH3 pathology. Furthermore, these findings open avenues for exploring astrocyte-targeted therapies for treating such devastating diseases.

## Methods

### Mutant Rats

#### Generation of mutant Pclo rat strains

Mutant *Pclo* rat strains harboring *Sleeping Beauty β-Geo* trap transposons were originally transmitted from a donor, recombinant rat spermatogonial stem cell library (Ivics et al. 2009). Recipient males were bred with wild-type females to produce a random panel of mutant rat strains enriched with gene trap in protein-coding genes (Izsvák et al. 2010). Rat use protocols were approved by the Institutional Animal Care and Use Committee (IACUC) at UT-Southwestern Medical Center in Dallas, as certified by the Association for Assessment and Accreditation of Laboratory Animal Care International (AALAC).

#### Genotyping of P0 - P2 pups

P0–P2 pups were genotyped via a PCR-based reaction. In brief, after dissection, the brain tissue was digested in a lysis buffer (100 mM Tris-HCl (pH 8.0), 100 mM NaCl, and 10 mg/ml proteinase K) at 55°C for 5 min. Subsequently, the proteinase K enzyme reaction was stopped, and the samples were incubated for 10 min at 99°C. Afterwards, the mixture was centrifuged at 14000 rpm for 2 min. Subsequently, PCR with Phire Tissue Direct PCR Master Mix (Thermo #F170S) was performed via specific primer combinations: F2: 3-gcaggaacacaaaccaacaa-5; R1: 3-tgacctttagccggaactgt-5; SBF2: 3-tcatcaaggaaaccctggac-5. The PCR protocol was as follows: 98°C for 5 min; 25 (98°C for 5 s, 55°C for 5 s, 72°C for 20 s); and 72°C for 1 min.

#### Pclo rat genotyping

Gene trap rats were genotyped via a PCR-based method. In brief, ear pieces taken from *Pclo* rats were digested overnight at 55°C in SNET buffer (400 mM NaCl, 1% SDS, 200 mM Tris (pH 8.0), 5 mM EDTA) containing 10 mg/ml proteinase K. Subsequently, the proteinase K enzyme reaction was stopped, and the samples were incubated for 10 min at 99°C. Afterwards, the mixture was centrifuged for 2 min at 14000 rpm. The supernatant was transferred into a fresh tube, and the DNA was precipitated through the addition of 100% isopropanol. The samples were subsequently centrifuged for 15 min at 13000 rpm. The precipitated DNA was washed once with 70% ethanol and centrifuged again for 5 min at 13000 rpm. Afterwards, the supernatant was discarded, and the DNA pellet was air dried and resuspended in H_2_O. PCR was performed on isolated DNA as previously described for P0 - P2 pub genotyping.

### Reverse Transcription PCR (RT‒PCR)

RT‒PCR was performed to assess gene expression levels. Total RNA was extracted from *Pclo^wt/wt^* and *Pclo^gt/gt^* primary cortical astrocytes as well as from cortical primary neurons via the SV Total RNA Isolation System Kit (Promega, Walldorf, Germany) following the manufacturer’s instructions. The RNA quality and concentration were determined via spectrophotometry. For reverse transcription followed by PCR amplification, the OneStep RT‒PCR Kit (Qiagen, Hilden, Germany) was used. Five to ten microliters of RNA was used to synthesize complementary DNA (cDNA) and amplify PCR fragments via OneStep RT‒PCR Enzyme Mix and 1 µl of dNTPs (10 mM of each dNTP) and 1.5 µl of gene-specific primers (PDZ-domain: fw 5’-AAATCACCAGAGACTCCAAGG-3’, rew 5’-AATCCAGCAGGTCAGCAAAG-3’; C2A-domain: fw 5’-TCCTTCAAGCAAGAAACCTAGTCC-3’, rew 5’-TGTCCTTCTCTTGTACTCAGC-3’) in a total volume of 25 µl. Reverse transcription and PCR amplification were carried out under the following conditions: reverse transcription at 50°C for 30 min, followed by PCR activation at 95°C for 15 min, followed by 38–40 cycles of denaturation at 94°C for 1 min, annealing at 51°C for 1 min, and extension at 72°C for 1 min, with a final extension step at 72°C for 10 min. PCR products were visualized via 1.5% agarose gel electrophoresis with Midori Green (Nippon Genetics Europe, Düren, Germany) staining.

### RNAseq analysis

#### RNA isolation

Total RNA was isolated from 50 mg of frozen cerebellar and brainstem tissue samples via mechanical homogenization in QIAzol Lysis Reagent (QIAGEN, Cat. No. 79306), followed by purification via the miRNeasy Mini Kit (QIAGEN, Cat. No. 217004) to ensure the retention of total RNA and other small RNA species, as per the manufacturer’s protocol. Elution of RNA was performed in 50 µL of RNase-free water, and the samples were subsequently stored at −80 °C until downstream processing. The quantification of RNA was conducted via a NanoDrop 2000 spectrophotometer (Thermo Fisher Scientific), and RNA integrity was assessed via the Agilent RNA ScreenTape assay on the TapeStation 4200 system (Agilent Technologies). All the RNA samples presented RNA integrity number (RIN) values greater than 9.5.

#### mRNA library preparation

Library preparation was performed using 1 µg of total RNA per sample with the TruSeq® Stranded mRNA Library Prep Kit (Illumina, Cat. No. 20020594) and TruSeq RNA Index Plate (Illumina, Cat. No. 20019792), following the manufacturer’s protocol for the mRNA HT low sample dual-index workflow. The final library amplification was carried out with 13 PCR cycles. Library quality and concentration were assessed via the D1000 ScreenTape assay on the Agilent 4200 TapeStation system. Equimolar pooling of libraries was performed, and sequencing was conducted on the Illumina NextSeq 550 platform via the NextSeq 500/550 High Output Kit (75-cycle, Illumina) to generate an average sequencing depth of ∼28 million reads per sample.

### GO analysis

GO analysis was conducted to explore the functional categorization of the differentially expressed genes (DEGs). The analysis was performed via G:Profiler (https://biit.cs.ut.ee/gprofiler/), a web-based tool that identifies enriched GO terms across biological processes, molecular functions, and cellular components. Enrichment analysis was carried out via the default statistical settings in G:Profiler, including correction for multiple testing with the false discovery rate (FDR) threshold set at <0.05. The results were further visualized as bar plots to highlight the most significant GO terms associated with the DEGs.

### Cell culture

#### Neuronal cell culture

Neuron cultures were prepared via the Banker culture protocol (Banker and Goslin 1988). Astrocytes derived from *Pclo^wt/wt^* and *Pclo^gt/gt^* P0-2 cortices or cerebella were plated into culture dishes 5–7 d before the addition of neurons (see below). Nitric acid-treated coverslips with paraffin dots were placed in separate culture dishes and covered with complete Neurobasal-A containing B-27 (Invitrogen, Thermo Fisher Scientific, Waltham, USA), 50 U/ml penicillin, 50 μg/ml streptomycin and 1x GlutaMAX (Invitrogen, Thermo Fisher Scientific, Waltham, USA). *Pclo^wt/wt^* and *Pclo^gt/gt^*cortices or cerebella were harvested from P0-2 brains in ice-cold HBSS (Gibco, Thermo Fisher Scientific, Waltham, USA) and incubated consecutively in 20 U/ml papain (Worthington, Lakewood, USA) for 30–45 min and 5 min in DMEM consisting of albumin (Sigma‒Aldrich, St. Louis, USA), a trypsin inhibitor (Sigma‒Aldrich, St. Louis, USA) and 5% FCS (Invitrogen, Thermo Fisher Scientific, Waltham, USA) at 37°C. Isolated cells were plated on coverslips with paraffin dots at a density of 60,000 cells/18 mm coverslips in 12-well plates and 30,000 cells/12 mm coverslips in 24-well plates in complete Neurobasal-A containing B27 (Invitrogen, Thermo Fisher Scientific, Waltham, USA), 50 U/ml penicillin, 50 μg/ml streptomycin and 1x GlutaMAX (Invitrogen, Thermo Fisher Scientific, Waltham, USA). After 2–3 h, the coverslips were flipped upside down and transferred to culture plates containing astrocytes in complete Neurobasal-A medium. Cultures were incubated at 37°C and 5% CO_2_ for 14–21 days before the experiments were started. On DIV 2-3, the cells were treated with 5 μM Ara-C (Sigma C6645).

ACM experiments, cortical astrocytes (60,000 cells/18 mm well) were seeded one week prior to neuron preparation. Two days before neuron preparation, seeded astrocytes were treated with FUDR (8 μM). Twenty-four hours before neuron culture, the astrocyte medium (DMEM) was replaced with Neurobasal-A containing B-27 (Invitrogen, Thermo Fisher Scientific, Waltham, USA), 50 U/ml penicillin, 50 μg/ml streptomycin and 1x GlutaMAX (Invitrogen, Thermo Fisher Scientific, Waltham, USA). Three hours after the neurons were seeded, the Neurobasal-A medium was replaced with warm astrocyte-conditioned medium (ACM). Half of the medium on growing neurons was subsequently replaced with ACM media every other day.

#### Astrocyte cell culture

For cortical or cerebellar astrocyte cell cultures, 1–2-week-old *Pclo^wt/wt^*and *Pclo^gt/gt^* astrocytes in T-75 flasks were vortexed for 1 minute and washed with PBS (Gibco #10010– 023). Subsequently, 5 mL of 0.05% trypsin-EDTA was added to each flask, followed by a 1-minute incubation. Afterwards, the supernatant was discarded, and the flasks were incubated at 37°C for another 7–8 minutes. The cells were detached by tapping the flask sides, resuspended in 8 mL of complete DMEM, transferred to Falcon tubes, counted, and seeded at the required densities of 30K, 60K, and 120 K in 24-, 12- or 6-well plates for neuronal culture and 30k in 12-well plates for astrocyte staining.

### Biochemistry and immunohistochemistry

#### Western blot

*Pclo^wt/wt^* and *Pclo^gt/gt^* brains (cortex, cerebellum) from P0–P2 pups as well as *Pclo*^wt/wt^ and *Pclo^gt/gt^* primary cortical or cerebellar astrocytes were lysed in lysis buffer (50 mM Tris-HCl, 150 mM NaCl, 5 mM EDTA, 1% Triton X-100, 0.5% deoxycholate, and protease inhibitor pH 7.5) and incubated on ice for 5 min. Subsequently, the samples were centrifuged at 13000 rpm for 10 min at 4°C. Afterwards, the supernatant was transferred into a fresh tube, and the protein concentration was determined via a BCA protein assay kit (Thermo Fisher Scientific, Waltham, Massachusetts, USA). The same protein amounts for the WT and GT samples were loaded and then separated via SDS‒PAGE (Bio-Rad, 4–15%, #4561084; 4–20%, #4561093DC; or 7.5%, #4561023) before being transferred onto nitrocellulose membranes (running buffer: 25 mM Tris, 190 mM glycine, 0.1% SDS, pH 8.3; transfer buffer: 25 mM Tris, 192 mM glycine, 1% SDS, 10% methanol for small proteins, and 7% methanol for larger proteins, pH 8.3). After transfer, the nitrocellulose membranes were blocked with 5% milk in TBST (20 mM Tris pH 7.5, 150 mM NaCl, 0.1% Tween 20) and incubated with primary antibodies in 3% milk in TBST o.n at 4°C. The following antibodies were used: Piccolo (1:500; rabbit; Synaptic Systems, Göttingen, Germany, Cat# 142 002, RRID:AB_887759 (AA 4439 to 4776); Tenascin-R (1:500, rabbit, Synaptic Systems, Göttingen, Germany, Cat# 217 008, RRID:AB_3083013); and Brevican (1:500, Thermo Fisher Scientific, Waltham, USA, Cat# MA5-45596, RRID:AB_2932050). The next day, the nitrocellulose membranes were washed 3 times for 10 min with TBST and incubated with HRP-labeled secondary antibodies for 1 h at RT (Thermo Fisher Scientific, Waltham, USA; dilution, 1:1000). The membranes were subsequently washed 3 times for 10 min with TBST, after which secondary antibody binding was detected with ECL Western Blotting Detection Reagents (Thermo Fisher Scientific, Waltham, Massachusetts, USA) and a Fusion FX7 image and analytics system (Vilber Lourmat).

#### Immunohistochemistry

Immunohistochemistry was performed on brain tissue from rats perfused transcardially with 4% PFA (Roth) dissolved in PB (80 mM NaH₂PO₄ (Roth), 20 mM NaH₂PO₄ (Bernd Kraft) (PFA). Following fixation, the brain was dissected from the head and further incubated in 4% PFA for 24 hrs at 4°C. The brain was subsequently transferred to 15% sucrose at 4°C for cryoprotection. After the tissue sank to the bottom of the tube, it was transferred into 30% sucrose and incubated again until it sank to the bottom. Afterwards, the brains were snap-frozen by submersion in 2-methylbutane (Roth, Karlsruhe, Germany), cooled to −60°C and stored at −20°C until use. The brains were cut parasagittally or coronally with a Leica Microsystems cryostat into either 20-µm-thick sections or 50 or 35-µm-thick sections, which were processed free-floating and mounted on superfrost slides (Thermo Fisher Scientific, from Roth, Karlsruhe, Germany). The slides were left to dry for a minimum of 1 h before being stored at 20°C. Free-floating sections were stored in antifreeze solution containing 30% ethylene glycol and 30% glycerol (Roth, Karlsruhe, Germany), 30% ddH₂O, 10% 0.244 M PO4 buffer, NaOH, and NaH2 PO4 (Roth, Karlsruhe, Germany) at −20°C.

Before staining, at least 4 slides (each containing two sections) from each animal were left to equilibrate at room temperature for 1 h. Sections were selected to encompass the range of the axis we were investigating. A hydrophobic barrier was created around the sections via a DAKO pen, and the sections were washed and permeabilized with TBS (20 mM Tris, pH 7.5; 150 mM NaCl (Roth, Karlsruhe, Germany) with 0.025% Triton X-100 (Roth, Karlsruhe, Germany) (TBST) for 3×5 min before being blocked with 10% normal goat serum (Sigma Millipore) with 1% BSA (Sigma Millipore) in TBS. The following primary antibodies were used: GFAP (1:1500-1:750); chicken; Millipore, Burlington, US, Cat# AB5541, RRID:AB_177521), Piccolo (1:200-1:400); rabbit; Synaptic Systems, Göttingen, Germany; Cat# 142002, RRID:AB_887759), and BCAN (1:100-1:250); rabbit; Proteintech, Illinois, US, Cat# 19017-1-AP, RRID:AB_10643526). The antibodies were diluted in TBS supplemented with 1% BSA and incubated overnight at 4°C. After 3×5 min of washing with TBST, different labeled secondary antibodies were used from Invitrogen (Thermo Fisher Scientific, dilution 1:1000), diluted again in 1% BSA in TBS antibody solution, and then incubated for 1 h at room temperature. The sections were then washed with TBS 2×10 min or, if desired, incubated in TBS with DAPI (Roth) for 30 min before being washed again for 2×10 min. The slides were coverslipped (24×50 mm coverslips, Menzel Gläser, Braunschweig, Germany) with Immuno-Mount (Shandon Thermo Scientific, Cheshire, UK) and sealed with clear nail polish once they hardened.

#### Immunocytochemistry (ICC)

Cortical neurons at DIV 14-16, as well as cortical and cerebellar astrocytes, were prepared for ICC. In brief, cells growing on coverslips were washed in PBS and subsequently fixed in 4% paraformaldehyde (PFA) for 5‒10 min at RT. After being washed 3 times in phosphate-buffered saline (PBS, Thermo Fisher Scientific, Waltham, USA), the cells were permeabilized with 0.1% Triton 100 in PBS (PBS-T). Afterwards, the cells were blocked in blocking solution (5% normal goat serum in PBS-T) for 30 min and incubated with primary antibodies in blocking solution overnight at 4°C. The following antibodies were used: Synaptophysin (1:1000; guinea pig; synaptic systems, Göttingen, Germany; Cat# 101 004, RRID:AB_1210382), PSD95 (1:500; mouse; Abcam, Cambridge, UK; Cat# ab2723, RRID:AB_303248), MAP2 (1:1000; chicken; Millipore, Darmstadt, Germany; Cat# AB5543, RRID:AB_571049), Piccolo (1:1000; rabbit; Synaptic Systems, Göttingen, Germany; Cat# 142002, RRID:AB_887759), GFAP (1:1000; chicken; Millipore, Burlington, US, Cat# AB5541, RRID:AB_177521), BCAN (1:200; rabbit; Proteintech, Illinois, US, Cat# 19017-1-AP, RRID:AB_10643526), TNR (1:200; rabbit; Synaptic Systems, Göttingen, germany; Cat# 217 008, RRID:AB_3083013), GM130 (1:300; mouse; BD Biosciences, Franklin Lakes, New Jersey, US, Cat# 610822, RRID:AB_398141), PRA1 (1:200; rabbit; Abcam, Cambridge, UK; Cat# ab213569), VGlut1 (1:1000; guinea pig; Synaptic Systems, Göttingen, germany; Cat# 135 304, RRID:AB_887878). Afterwards cells were washed 3 times with PBS-T and incubated with secondary antibodies for 1 h at RT. Differently labeled secondary antibodies were obtained from Invitrogen (Thermo Fisher Scientific, Waltham, USA; dilution, 1:1000). After washing, the coverslips were mounted in ProLong Diamond Antifade Mountant (Thermo Fisher Scientific, Waltham, USA) and stored at 4°C until imaging.

### Electrophysiology

Whole-cell voltage‒clamp experiments were performed via one channel of a MultiClamp 700B amplifier (Molecular Devices, Sunnyvale, USA) under the control of a Digidata 1440A Digitizer (Molecular Devices, Sunnyvale, USA) and pCLAMP Software (Molecular Devices, Sunnyvale, USA). Neurons in cortical mass cell cultures were recorded at DIV 14–21. The cortical neurons were clamped at −70 mV, the data were sampled at 10 kHz, and Bessel filtered at 3 kHz. The series resistance was typically under 10 MΩ. The series resistance was compensated by at least 70%. The extracellular solutions for all the experiments, unless otherwise indicated, contained the following (in mM): 140 NaCl, 2.4 KCl, 10 HEPES, 10 glucose, 4 MgCl_2_, and 2 CaCl_2_ (pH 7.4). The intracellular mixture contained the following (in mM): 126 KCl, 17.8 HEPES, 1 EGTA, 0.6 MgCl_2_, 4 MgATP, 0.3 Na_2_GTP, 12 creatine phosphate, and phosphocreatine kinase (50 U/mL) (300 mOsm, pH 7.4). All reagents were purchased from Carl Roth GMBH (Essen, Germany), with the exception of Na_2_ATP, sodium Na_2_GTP, creatine-phosphokinase (Sigma‒Aldrich, St. Louis, MO), and phosphocreatine (EMD Millipore Chemicals, Billerica, USA).

All recordings were performed in the presence of TTX (0.5 μM, Tocris) to block the propagation of action potentials. A sucrose response, representing the readily releasable pool (RRP) of synaptic vesicles, was evoked by a 5-second application of hypertonic sucrose (500 mM in extracellular solution). The RRP size was estimated by integrating the charge of the sucrose-evoked current, using the steady-state current as the baseline. To isolate excitatory and inhibitory RRP components, experiments were repeated in the presence of either 30 µM bicuculline (GABA receptor antagonist; Tocris) to measure excitatory RRP or 3 µM NBQX (AMPA receptor antagonist) and 10 µM AP5 (NMDA receptor antagonist) to measure inhibitory RRP. Miniature excitatory (mEPSCs) and miniature inhibitory (mIPSCs) postsynaptic currents were recorded in si× 10-second sweeps. mEPSCs were recorded in the presence of 100 nM TTX and 30 µM bicuculline, while mIPSCs were recorded with 100 nM TTX, 3 µM NBQX, and 10 µM AP5. Electrophysiological recordings were analyzed via Axograph X (Axograph, Berkeley, USA), Excel (Microsoft, Redmond, USA), and Prism software (GraphPad, La Jolla, USA). The recorded traces were filtered at 1 kHz to reduce electrical noise. The initial sweep of each recording was discarded to ensure drug equilibration. Miniature events were detected via a template-based algorithm in Axograph X, with a detection threshold set at three times the standard deviation of the baseline noise. mEPSCs were identified as events exceeding a 10 pA amplitude threshold, with a rise time of 0.15–1.5 ms and a half-width of 0.5–5 ms. mIPSCs were defined as events exceeding a 10 pA threshold, with a rise time of 0.2–1.5 ms and a half-width of 1–30 ms.

### Image acquisition and processing

#### Imaging

Images for immunocytochemical and histochemical staining were acquired on a spinning disc confocal microscope (Carl Zeiss Axio Observer. Z1 with an Andor spinning disc and cobolt, omricron, and i-beam laser (405, 490, 562 and 642 nm wavelengths) using a 40 (1.3 NA) and 63 (1.4 NA) Plan-Apochromat oil objective and an iXon ultra (Andor) camera controlled by iQ software (Andor).

#### Image analyses

For image processing and analysis, ImageJ/FIJI software was used (Schindelin et al. 2012).

#### Cargo distribution in astrocytes

The intensity of cargo molecules (BCAN, TNR) along the secretory pathway from the nucleus toward the plasma membrane in cortical and cerebellar primary astrocytes was measured via a Python script. A circle was drawn around the nucleus (DAPI); afterwards, a line was drawn at a 45° angle from the center of the nucleus to the plasma membrane. The line was subsequently divided into 10 equal boxes/bins, and the mean intensity in each box was measured. The experimenter was blinded to all the genotypes.

#### Synapse density

To determine synapse density, the number of Synaptophysin-PSD95 double-positive puncta along MAP2-positive primary dendrites was determined. Puncta per unit length of dendrite were counted manually. The experimenter was blinded to all the genotypes.

#### BCAN intensity in brain slices

Astrocytes were identified by their expression of glial fibrillary acidic protein (GFAP). GFAP-positive regions were detected through a standardized thresholding method, which ensured consistent delineation of astrocytic structures across images. Following thresholding, the “Analyze Particles” tool in Fiji (Schindelin et al. 2012) was used to isolate and quantify the identified astrocytic regions. BCANcan intensity was subsequently measured within these regions of interest (ROIs). This was achieved by overlaying the identified astrocytic masks onto the corresponding BCAN fluorescence channel. The IntDen values for BCAN were extracted from each astrocytic ROI, enabling quantitative assessment of BCAN expression in GFAP-positive astrocytes.

To measure the intensity of BCAN in areas outside astrocytes, GFAP-positive astrocytes were first identified as described previously. The defined astrocytic regions were subsequently used to create an inverse mask, representing the area outside the astrocytes. The inverse mask was then applied to the BCAN fluorescence channel, allowing for the exclusion of astrocytic regions from the analysis. BCAN intensity was then quantified within the nonastrocytic areas by measuring the IntDen fluorescence of BCAN in the inverse mask. The experimenter was blinded to all the genotypes.

#### GM130 analysis

The GM130 surface area was measured through a standardized thresholding method, which ensured consistent delineation of the GM130 structures across the images. Following thresholding, the “Analyze Particles” tool in Fiji (Schindelin et al. 2012) was used to quantify the area of the GM130-positive regions. The experimenter was blinded to all the genotypes.

#### Piccolo intensity in primary astrocytes

Pclo intensity at the Golgi (GM130 positive) was measured in *Pclo^wt/wt^* and *Pclo^gt/gt^*. Therefore in the GM130 channel the GM130-positive Golgi ROI was thresholded and selected using the “Analyze Particles” tool in Fiji (Schindelin et al. 2012). Subsequently the defined ROI was applied to the Pclo channel and Pclo IntDen was measured within this area.

#### Pra1 intensity in primary astrocytes

Pra1 intensity at the Golgi (GM130 positive) was measured in *Pclo^wt/wt^* and *Pclo^gt/gt^*. Therefore in the GM130 channel the GM130-positive Golgi ROI was thresholded and selected using the “Analyze Particles” tool in Fiji (Schindelin et al. 2012). Subsequently the defined ROI was applied to the Pra1 channel and Pra1 RAWIntDen was measured within this area.

#### BCAN intensity at VGlut1 positive presynaptic terminals

BCAN intensity at VGlut1 puncta was measured in neuron cultures grown together with *Pclo^wt/wt^* and *Pclo^gt/gt^*. In the VGlut1 channel VGlut1 positive ROIs were threshold and selected using the “Analyze Particles” tool in Fiji (Schindelin et al. 2012). Subsequently the defined ROIs were applied to the BCAN channel and BCAN IntDen was measured within these areas.

#### Western blot analysis

The lanes containing protein bands were defined via the “Analyze Gels” tool in Fiji (Schindelin et al. 2012), which involves selecting and aligning the lanes for proper comparison. The intensity profiles of the lanes were subsequently generated, allowing for the identification and quantification of individual protein bands as peaks in the profile. The area under each peak, corresponding to the band intensity, was quantified and recorded for downstream statistical analysis.

### Statistical analysis

Statistical significance was assessed via GraphPad analysis. The data were tested for normality via the following tests: the Anderson‒Darling test, the D’Agostino‒Pearson omnibus normality test, the Shapiro‒Wilk normality test, and the Komogorov‒Smirnov normality test with the Dallal‒Wilkinson‒ Lillie test for p values. For normally distributed data, Student’s t test or one-way ANOVA was used to determine significance, and the values are shown as the means± SEMs. For nonparametric data, the Mann‒Whitney test was used to determine significance, and the values are shown as the median ± interquartile range (IQR). For all the statistical tests, the 0.05 confidence level was considered statistically significant. In all figures, * denotes p < 0.05, ** denotes p < 0.01, *** denotes p < 0.001 and **** denotes p < 0.0001.

## Results

### Loss of Piccolo function led to differentially expressed genes (DEGs) in the cerebellum and brainstem of *Pclo^gt/gt^* rats during postnatal development and in adulthood

Piccolo is a large (>420 kDa) presynaptic scaffolding protein (S. D. Fenster et al. 2000); its loss of function in rats leads to a complex phenotype consisting of multiple cellular and tissue malfunctions (Garner and Ackermann 2023; A. Medrano et al. 2018). Until now, how the loss of a synaptic protein, like Piccolo, can cause such a pleiotropic effect on phenotypes and symptoms has not been well understood. To better understand these changes, this study aimed to identify altered cellular/signaling pathways in the *Pclo^gt/gt^*rat brain. Analyzing gene expression profiles is a well-established approach to identify processes that may be disrupted upon gene mutation. Therefore, we first performed RNA-seq analysis on brain regions that are specifically affected in *Pclo^gt/gt^* animals, the cerebellum and the brainstem. Additionally, we analyzed two developmental stages—early postnatal development (P5), when GC proliferation is initiated in the cerebellum, and young adulthood (P25), when the cerebellar microcircuitry has been established but is still immature—to investigate previously reported changes in the number of GCs and MF formation (Figure 1). During early postnatal development (P5), only a small subset of genes is differentially expressed in *Pclo^gt/gt^*compared with the *Pclo^wt/wt^* cerebellum and brainstem tissue (Figure 1 A). Here, we define differentially expressed genes (DEGs) as those whose gene expression is changed by at least 0.5 log2-fold (log2FD) with an adjusted p value smaller than 0.05 compared with that of *Pclo^wt/wt^*cerebellum and brainstem tissue. At P5, we identified 12 DEGs in the *Pclo^gt/gt^*brainstem and 39 DEGs in the cerebellum *Pclo^gt/gt^* tissue (Figure 1 A, C, D, P5). Intriguingly, this number significantly increased in P25 rats. Here, we identified 233 DEGs in the *Pclo^gt/gt^* brainstem dataset and 1563 DEGs in the *Pclo^gt/gt^* cerebellum dataset (Figure 1 A, E, F, P25), indicating that Piccolo loss-of-function phenotypes progress over time during postnatal development. The expression of affected genes changed in both directions, with some DEGs being upregulated (red) and others being downregulated (blue) (Figure 1 C - F). During early postnatal development (P5), most DEGs were downregulated (Figure 1 C and D, cerebellum (CB): 31; brainstem (BS): 7); however, this ratio changed by P25, when the expression of most DEGs was upregulated (red) (Figure 1 E and F, BS: 169; CB: 1124), and a small subset was downregulated (blue) (Figure 1 E and F, BS: 64; CB: 439). Since a SNP mutation in Piccolo has been associated with PCH3, we further investigated whether genes associated with other PCH subtypes are also affected in our RNA-seq dataset (Figure 1 B, adapted from (van Dijk et al. 2018)). We did not find matching genes but rather three genes from the same protein family (Figure 1 B (red)). Among those genes, two are downregulated (*SLC25A25* and *TBC1D4*), and one is upregulated (*CHMP4C*) (Figure 1 B), indicating that known genes implicated in the progression of other PCH subtypes most likely do not contribute to the *Pclo^gt/gt^*phenotype.

**Fig. 1.**
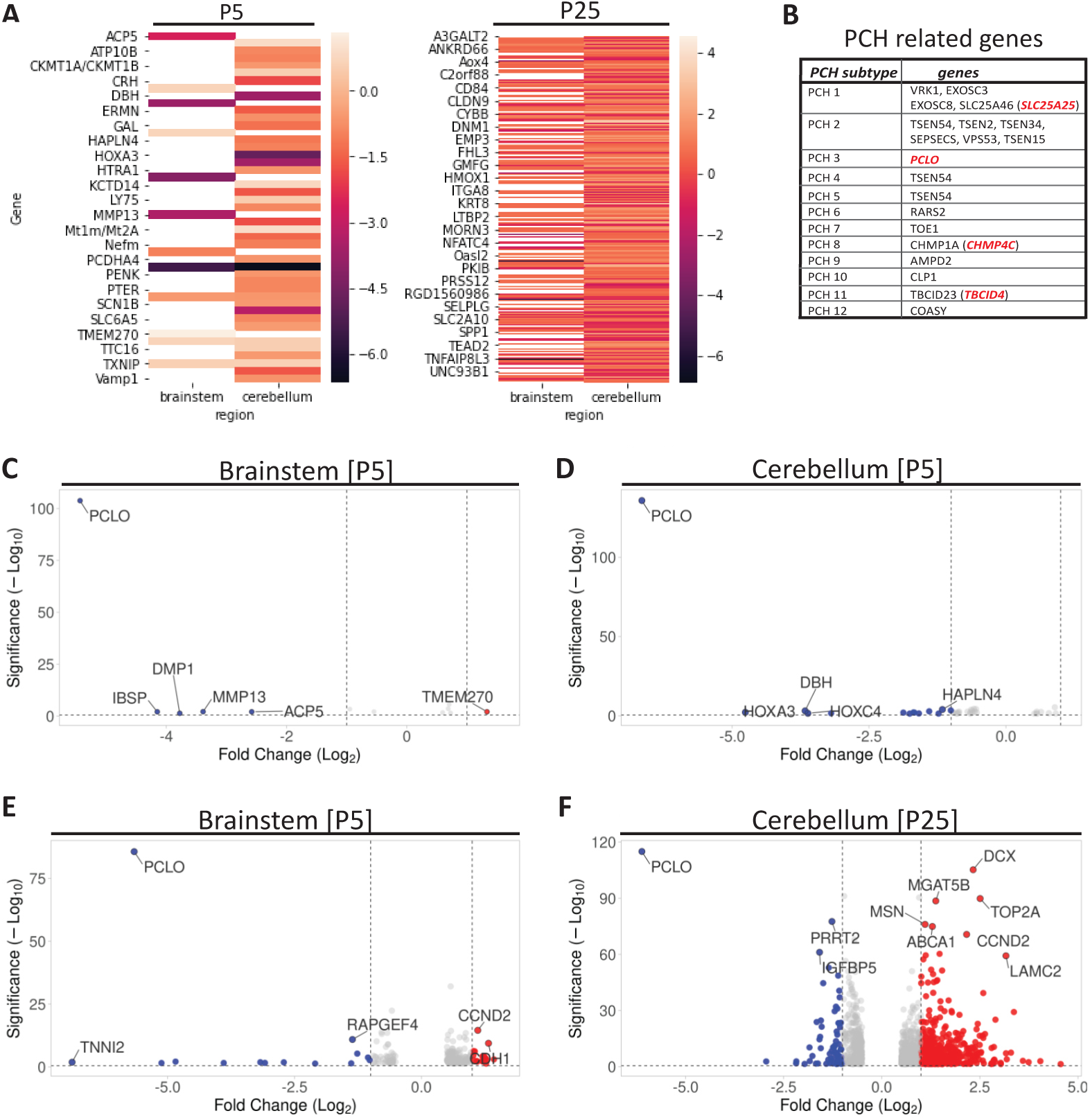
Most genes were differentially expressed in P25 *Pclo^gt/gt^* cerebellum tissue. **A**, Heatmap showing the expression of DEGs in the brainstem and cerebellum at P5 and P25. The majority of DEGs were observed in P25 cerebellum tissue, with both upregulated and downregulated genes. In contrast, very few genes were differentially expressed in P5 brainstem tissue. **B**, Table listing known genes associated with various PCH subtypes. In the *Pclo^gt/gt^*RNA-seq dataset, related genes are highlighted in red. Among these genes, *SLC25A25*, which is associated with *SLC25A46*, a gene linked to PCH1, was identified. Similarly, *CHMP4C*, related to *CHMP1A* (linked to PCH8), and *TBC1D4*, related to *TBC1D23* (linked to PCH11), were also identified. **C**, Volcanoblot displaying DEGs found in the *Pclo^gt/gt^* brainstem P5 RNA-seq dataset (blue: downregulated; red: upregulated; settings: −1 to 1-fold change; significance threshold 0.5; Manhattan distance ranking). In total, 12 DEGs were identified, most of which were downregulated. **D**, Volcanoblot displaying DEGs found in the *Pclo^gt/gt^* cerebellum P5 RNA-seq dataset (blue: downregulated; red: upregulated; settings: −1 to 1-fold change; significance threshold 0.5; Manhattan distance ranking). In total, 39 DEGs were found, most of which were downregulated. **E**, Volcanoblot displaying DEGs found in the *Pclo^gt/gt^*brainstem P25 RNA-seq dataset (blue: downregulated; red: upregulated; settings: −1 to 1-fold change; significance threshold 0.5; Manhattan distance ranking). In total, 233 DEGs were found, which were up- or downregulated. **F**, Volcanoblot displaying DEGs found in the *Pclo^gt/gt^* cerebellum P25 RNA-seq dataset (blue: downregulated; red: upregulated; settings: −1 to 1-fold change; significance threshold 0.5; Manhattan distance ranking). In total, 1561 DEGs were found, which were up- or downregulated.

In summary, our initial RNA-seq dataset revealed that Piccolo loss of function has a limited effect on gene expression in early postnatal *Pclo^gt/gt^* cerebellum and brainstem tissues; however, the magnitude of alterations increases over time.

### DEGs in the cerebellum and brainstem of adult *Pclo^gt/gt^* rats are enriched in gene ontology (GO) terms associated with signaling and the extracellular matrix

As we only observed a small subset of DEGs in *Pclo^gt/gt^*brainstem and cerebellum tissue at P5, we performed all subsequent analyses on the P25 *Pclo^gt/gt^* RNA-seq dataset (brainstem, cerebellum). Here, we performed a GO enrichment analysis via the open-source software g:Profiler to investigate the functional annotation of the identified DEGs. We analyzed the enrichment of GO terms in three categories: biological process (BP), molecular function (MOF), and cellular component (CC). The analysis revealed few enriched molecular functions for the cerebellum RNA-seq dataset, including “protein binding”, “signal receptor binding” and “extracellular matrix binding” (Figure 2 B, MF enrichment scores: 37.53, 7.04, and 4.32, respectively). Moreover, the biological processes “regulation of cell communication”, “signaling” and “ECM organization” (Figure 2 B, BP enrichment score: 30.25, 22.67, 14.38) and the cell components “cell periphery”, “plasma membrane” and “ECM” were enriched (Figure 2 B, CC enrichment score: 73.66, 56.52, 16.27). Compared with the cerebellum, the molecular function “protein binding” was also enriched in the brainstem RNA-seq dataset (Figure 2 A, MF enrichment score: 3.81). Similarly, in the brainstem, the biological processes “regulation of cell communication”, “ECM organization” and “cell adhesion” (Figure 2 A, BP enrichment score: 5.58, 4.92, 6.41) and the cell components “cell periphery”, “plasma membrane” and “ECM” were enriched (Figure 2 A, CC enrichment score: 13.99, 9.23, 8.19).

**Fig. 2.**
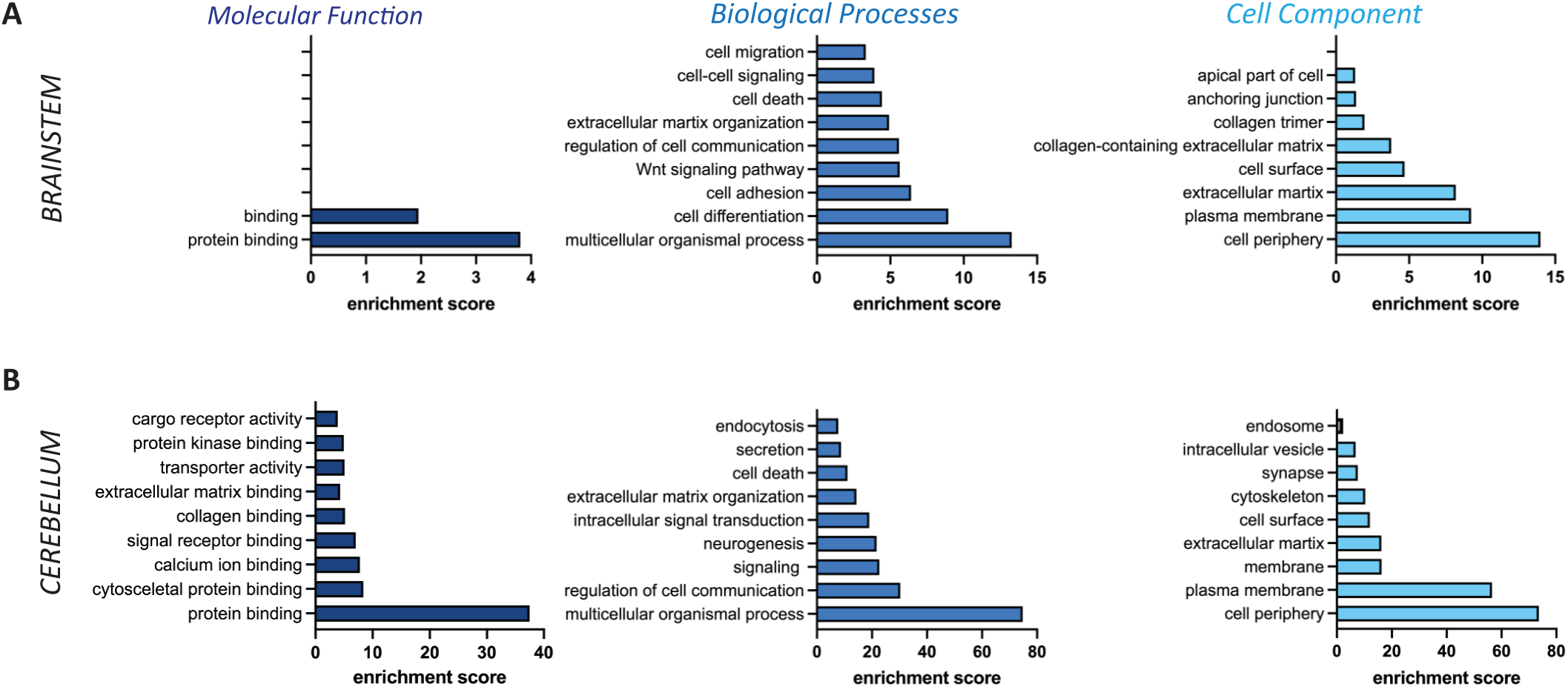
Many DEGs were enriched in GO categories related to cell‒cell communication, the ECM and signaling. **A**, GO enrichment analysis of brainstem P25 DEGs. The analysis revealed only two enriched molecular functions: “binding” (1.95) and “protein binding” (3.80). Several biological processes were enriched, including “regulation of cell communication” (5.58), “ECM organization” (4.92) and “cell adhesion” (6.41). Among the cell components “cell periphery” (13.99), “plasma membrane” (9.23) and “extracellular matrix” (8.18) were enriched. **B**, GO enrichment analysis of cerebellar P25 DEGs. The analysis revealed a few enriched molecular functions, including “protein binding” (37.53), “signal receptor binding” (7.04) and “ECM binding”. Several enriched biological processes included “regulation of cell communication” (30.25), “signaling” (22.67) and “ECM organization” (14.38). Among the cell components, “cell periphery” (73.65), “plasma membrane” (56.52) and “ECM” (16.27) were enriched.

Together, these findings reveal a comparable enrichment of DEGs in brainstem and cerebellum tissues and suggest that changes in gene expression are general in nature and affect signaling pathways across different tissues, some of which are involved in cell communication, signaling and the ECM. These changes offer etiological insights into the symptoms observed in children with PCH3.

### Secretion of the ECM component BCAN is impaired from cerebellar astrocytes in *Pclo^gt/gt^* brain sections

In our *Pclo^gt/gt^* PCH3 rat model, the main subcellular phenotype is smaller and disorganized cerebellar mossy fiber (MF) synapses, which are formed between pontine axons and granule cells in the granule cell layer (GCL) of the cerebellum (Falck et al. 2020). This finding points to defects in synaptogenesis and/or synapse maturation. Interestingly, our GO analysis (Figure 2) highlighted the ECM in all categories. The ECM is an important structural and functional component of the brain that regulates the formation and function of synapses (Barros, Franco, and Müller 2011). Therefore, we aimed to investigate a potential link between the ECM, altered synapses and other PCH3 phenotypes in *Pclo^gt/gt^*rats. First, we examined ECM expression in *Pclo^wt/wt^* and *Pclo^gt/gt^* brain sections. Specifically, we initially analyzed the levels and distribution of the ECM component BCAN. We choose it because it is highly expressed in the cerebellum and associated with astrocytes ensheathing cerebellar glomeruli (Yamada et al. 1997), which are those synaptic structures that are altered in our Pclo rat model (Falck et al. 2020). In addition, BCAN is present at the perisynaptic matrix and known to mediate synapse density as well as plasticity (Nakajo et al. 2025; Frischknecht and Seidenbecher 2012). Therefore, we stained P25 in cerebellar brain sections via antibodies against BCAN and the astrocytic intermediate filament protein glial fibrillary acidic protein (GFAP) (Figure 3 A). Here, BCAN is present not only in GFAP-positive astrocytes in the *Pclo^wt/wt^* and *Pclo^gt/gt^* cerebellar granule cell layers but also in the surrounding extracellular tissue, as BCAN is secreted from astrocytes (Yamada et al. 1997; John et al. 2006) (Figure 3 A). Therefore, we quantified the amount (fluorescence intensity) of BCAN inside and outside astrocytes (Figure 3 B, C). Inside astrocytes, we observed similar BCAN levels between *Pclo^wt/wt^* and *Pclo^gt/gt^* cerebellar brain sections (Figure 3 A, B; *Pclo^wt/wt^*mean± SEM= 1± 0.082, n=3 animals; *Pclo^gt/gt^* mean± SEM= 1.151± 0.082, n=2 animals, p=0.082 t test). In contrast, the level of BCAN secreted from astrocytes was significantly decreased in *Pclo^gt/gt^* cerebellar brain sections (Figure 3 A, C; *Pclo^wt/wt^*mean± SEM= 1± 0.069, n=3 animals; *Pclo^gt/gt^* mean± SEM= 0.8315± 0.069, n=2 animals, p=0.018, t test), which led us to hypothesize that the secretion of BCAN from astrocytes is altered in *Pclo^gt/gt^* brains due to Piccolo loss of function.

**Fig. 3.**
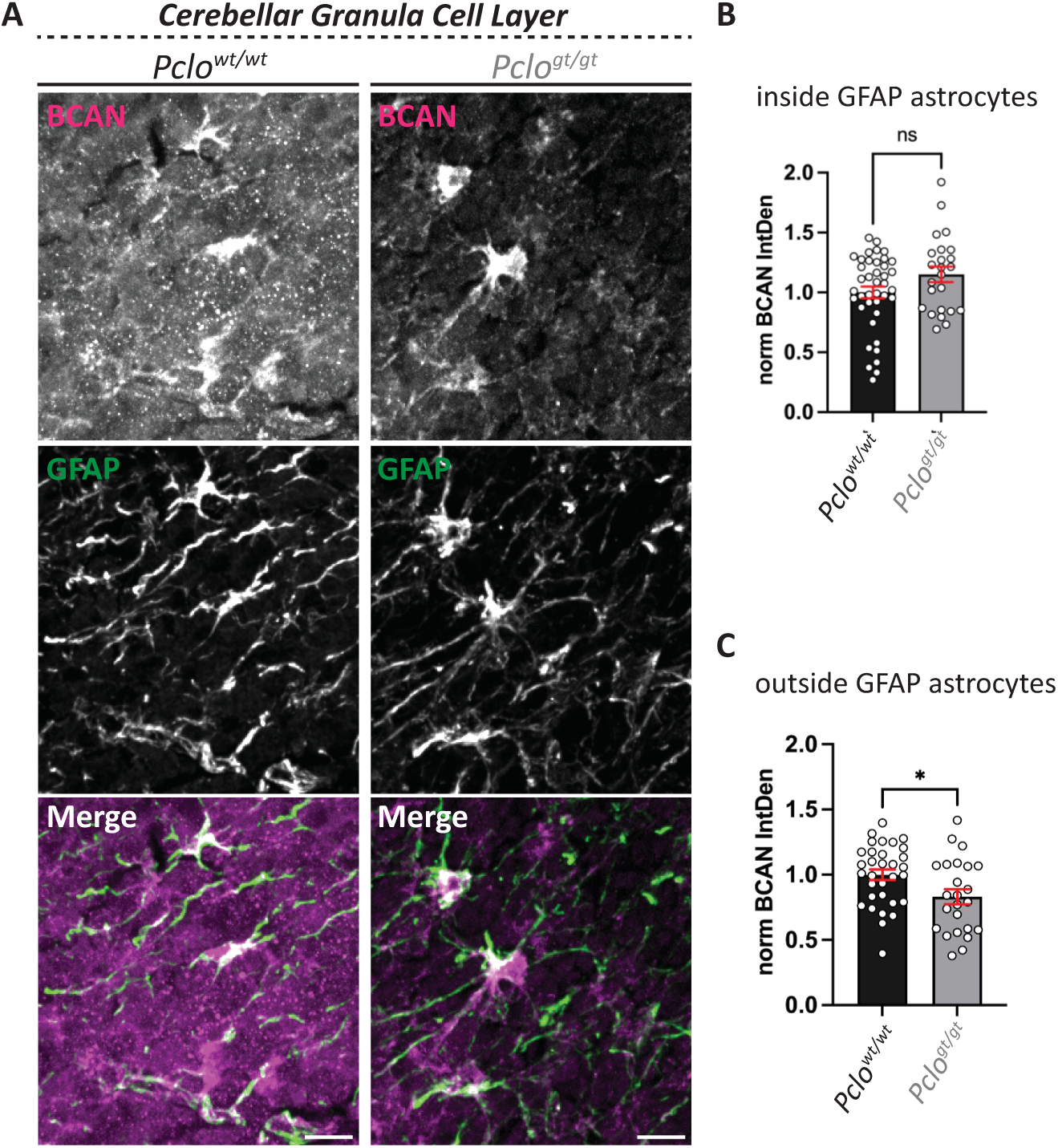
Secretion of BCAN from astrocytes in *Pclo^gt/gt^* cerebellar brain sections is impaired. **A,** Immunohistological staining for BCAN was performed on cerebellar brain sections from *Pclo^wt/wt^* and *Pclo^gt/gt^*rats. In both genotypes, BCAN is localized in GFAP-positive astrocytes, from which it is secreted. In *Pclo^wt/wt^* slices, BCAN is also found in the surrounding tissue, indicating effective secretion. In contrast, in *Pclo^gt/gt^* sections, significantly less BCAN was detected outside astrocytes, suggesting impaired secretion of BCAN from astrocytes. **B**, Quantification of a. BCAN intensity is slightly greater inside astrocytes in *Pclo^gt/gt^* cerebellar sections than in *Pclo^wt/wt^* cerebellar slices; however, this difference is not significant (*Pclo^wt/wt^*: mean± SEM= 1± 0.05, n=40 astrocytes, 3 independent animals; *Pclo^gt/gt^*: mean± SEM= 1.151± 0.06, 2 independent animals; p=0.082, t test). **C,** Quantification of a. BCAN intensity is significantly lower outside astrocytes in *Pclo^gt/gt^* slices than in *Pclo^wt/wt^* slices (*Pclo^wt/wt^*: mean± SEM= 1± 0.04, 3 independent animals; *Pclo^gt/gt^*: mean± SEM= 0,831± 0.06, 2 independent animals; p=0.018, t test). Scale bar, 10 μm.

### Piccolo is present in brain astrocytes

This finding is exciting and surprising, as it indicates an additional role for Piccolo in astrocytes. In fact, Song and colleagues demonstrated Piccolo in GFAP-positive astrocytes in brain sections of mice exposed to an enriched environment (Song et al. 2018). To confirm the presence of Piccolo in astrocytes, we initially conducted RT‒PCR analysis. We used primers targeting the housekeeping gene *HPRT1* as RNA quality control (Figure S1 C), in addition we used primers designed to amplify regions of Piccolo’s PDZ and C2A domains (Figure 4 A and B). When primers targeting the PDZ domain were used, a double band (140, 210 bp) was amplified from *Pclo^wt/wt^* mRNA isolated from primary neurons, indicating that Piccolo is expressed as two distinct PDZ domain structures/isoforms in neurons (Figure 4 A, neurons). In contrast, no bands were amplified from the corresponding *Pclo^gt/gt^*mRNAs, suggesting that both Piccolo isoforms are lost in *Pclo* gene trap neurons (Figure 4 A, neurons). Interestingly, the smaller band, but not the higher band, was also amplified from *Pclo^wt/wt^*mRNA isolated from primary cortical rat astrocytes, supporting the hypothesis that distinct Piccolo isoforms are expressed in astrocytes, which only harbor the smaller PDZ domain structure. These data suggest that astrocytes possess a Piccolo protein pool, which is distinct from that in neurons (Figure 4 A, cortical astrocytes). In contrast, a weaker small band and a stronger larger band, similar to that observed in neurons, can be amplified from *Pclo^gt/gt^* mRNA isolated from cortical astrocytes (Figure 4 A, cortical astrocytes). These findings further indicate that the occurrence of Piccolo isoforms, which affect the PDZ domain, is altered in *Pclo^gt/gt^*gene trap astrocytes.

**Fig. 4.**
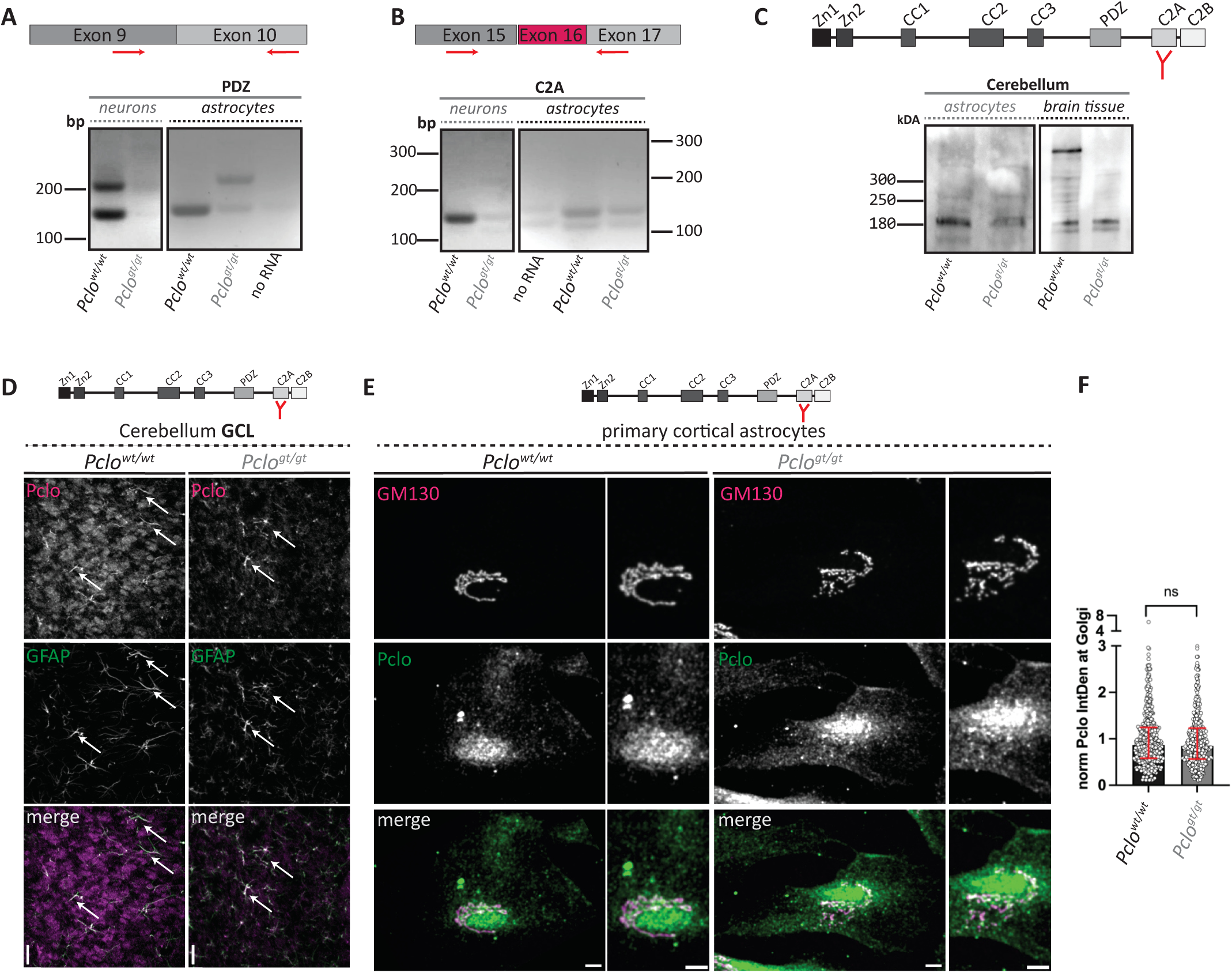
Piccolo is present in primary astrocytes. **A,** RT‒PCR using primers for Piccolo’s PDZ domain. A double band was amplified from *Pclo^wt/wt^* cortical neuron RNA but not from Pclo*^gt/gt^*. A single band was amplified from *Pclo^wt/wt^*cortical astrocyte RNA. In contrast, a double band similar to that in *Pclo^wt/wt^*cortical neurons was amplified from *Pclo^gt/gt^*cortical astrocyte RNA. **B,** RT‒PCR using primers for Piccolo’s C2A domain. A positive band was amplified from *Pclo^wt/wt^* cortical neuron RNA but not from *Pclo^gt/gt^*. In *Pclo^wt/wt^* cortical astrocytes, a double band was amplified. In contrast, weaker bands were amplified from *Pclo^gt/gt^* cortical astrocytes. **C,** WB analysis. Full-length Piccolo is present in *Pclo^wt/wt^* but is absent in *Pclo^gt/gt^* brain tissue. Similarly, full-length Piccolo is absent in both *Pclo^wt/wt^* and *Pclo^gt/gt^*cortical astrocyte lysates; however, a smaller isoform (<180 kDa) is present in *Pclo^wt/wt^* and *Pclo^gt/gt^* cortical astrocytes. **D,** Immunohistological staining. Using an antibody targeting the C2A region of Piccolo, positive synaptic staining was observed in *Pclo^wt/wt^* but not in *Pclo^gt/gt^* cerebellar sections. Intriguingly, Piccolo was also detected in GFAP-positive *Pclo^wt/wt^*and *Pclo^gt/gt^* astrocytes. **E**, Immunocytochemical staining of primary cortical astrocytes. An antibody targeting the C2A region of Piccolo was used. Positive staining for Piccolo is detected in cortical astrocytes, which colocalize with GM130. There was no significant difference in Piccolo levels between *Pclo^wt/wt^* and *Pclo^gt/gt^* astrocytes at the GM130-positive Golgi apparatus. **F,** Quantification of e. There was no difference in Piccolo levels at the Golgi apparatus between cortical *Pclo^wt/wt^* and *Pclo^gt/gt^* astrocytes (*Pclo^wt/wt^*: median (IQR)= 0.865 (0.665), 3 independent experiments; *Pclo^gt/gt^*: median (IQR)= 0.847 (0.660), 3 independent experiments; p=0.7891, Mann‒Whitney test). Scale bars, 10 μm.

Probing the presence of the C2A domain sequence, we amplified a single band from *Pclo^wt/wt^* mRNA isolated from primary cortical neurons. Notably, this region/band was not amplified from *Pclo^gt/gt^*mRNA isolated from primary cortical neurons (Figure 4 B, cortical neurons). However, this band, along with an additional smaller and weaker band, could also be amplified from astrocytic *Pclo^wt/wt^* mRNAs (Figure 4 B, cortical astrocytes). The smaller band is most likely due to alternative splicing of exon 16 (Figure 4 B), again indicating that Piccolo isoforms with different domain structures exist in astrocytes. Intriguingly, the ratio between the isoforms seems to shift upon transposon mutagenesis in *Pclo^gt/gt^* astrocytes (Figure 4 B, cortical astrocytes).

Together, our RT‒PCR analysis results suggest that Piccolo isoforms, which contain both PDZ and C2A domains, are expressed in *Pclo^wt/wt^*astrocytes and undergo differential splicing in *Pclo^gt/gt^* astrocytes due to transposon mutagenesis. Next, we investigated whether the Piccolo protein is also present in astrocytes. Therefore, we performed Western blot analysis on total brain lysates and primary astrocyte lysates (Figure 4 C). Here, we identified a band representing Piccolo full-length protein (560 kDa) in *Pclo^wt/wt^* cerebellar brain tissue via an antibody raised against the Piccolo C2A domain. In contrast, this band was absent in *Pclo^gt/gt^* cerebellar brain tissue. Intriguingly, the Piccolo full-length band was absent in *Pclo^wt/wt^* and *Pclo^gt/gt^*primary astrocytes, although a smaller band at 170 kDa was present in both samples (Figure 4 C), which could represent astrocyte-specific Piccolo isoforms containing the C2A domain. Notably, this band is also present in total brain tissue, which represents a mixture of all brain cell types, including neurons and astrocytes. We investigated the presence and localization of Piccolo in cerebellar brain sections and primary cortical astrocytes. Immunohistological staining of cerebellar brain sections from *Pclo^gt/gt^* and *Pclo^wt/wt^* rats via an antibody raised against the C2A domain of Piccolo revealed, as described previously (Ackermann et al. 2019; Falck et al. 2020), synaptic staining for Piccolo in *Pclo^wt/wt^* cerebellar sections, which was absent in *Pclo^gt/gt^* sections (Figure 4 D). Interestingly, we also observed additional labeling for Piccolo in GFAP-positive astrocytes, although it was obscured by synaptic staining (Figure 4 D, *Pclo^wt/wt^*; arrow). While synaptic staining is absent in *Pclo^gt/gt^* sections, astrocytic staining remains intact (Figure 4 D, *Pclo^gt/gt^*; arrow), supporting the hypothesis that various Piccolo isoforms still exist after transposon mutagenesis in the brain and localize differently to synapses and astrocytes. Because we hypothesized that there was a change in the secretion properties of *Pclo^gt/gt^* astrocytes, we investigated whether Piccolo was associated with the protein trafficking pathway. Similarly, we also observed Piccolo-positive staining in primary *Pclo^wt/wt^* and *Pclo^gt/gt^*astrocytes, which partially colocalized with the Golgi apparatus marked by GM130, equally between *Pclo^wt/wt^* and *Pclo^gt/gt^* (Figure 4 E and F, *Pclo^wt/wt^*: median (IQR)= 0.865 (0.665), n=3 independent experiments; *Pclo^gt/gt^*: median (IQR)= 0.847 (0.660), n=3 independent experiments; p=0.7891, Mann‒Whitney test).

Taken together, these data indicate that Piccolo isoforms are present in astrocytes and become alternatively spliced upon transposon mutagenesis, leading to an altered ratio of Piccolo isoforms in *Pclo^gt/gt^* astrocytes.

### The subcellular distributions of BCAN and TNR are altered in *Pclo^gt/gt^* primary astrocytes

The initially observed reduction in BCAN in the intercellular tissue of *Pclo^gt/gt^* cerebellar brain sections suggests that the secretion of this ECM component from astrocytes is impaired due to Piccolo gene trap mutation. Thus, we next investigated the subcellular localization/distribution of BCAN and another secreted ECM component, in primary cortical astrocytes prepared from *Pclo^wt/wt^* and *Pclo^gt/gt^*rat brains (Figure 5 A, B). TNR was chosen as it is known to interact with BCAN, to form a network which subsequently helps to stabilize the ECM (Hagihara et al. 1999). Thus, we analyzed the distribution of BCAN and TNR throughout the cell from the endoplasmic reticulum (ER)/Golgi toward the cell periphery. To do so, we laid a fictitious line from the nucleus toward the cell periphery and subdivided it into 10 bins. Afterwards, we measured the fluorescence intensity of the distinct cargo molecules (BCAN/TNR) in these defined bins (1=nucleus – 9=plasma membrane) (Figure 5 A - D). Interestingly, the distribution of the TNR is altered toward the cell periphery in primary cortical *Pclo^gt/gt^* astrocytes (Figure 5 B and D). Compared with that in *Pclo^wt/wt^* astrocytes, the fluorescence intensity in *Pclo^gt/gt^* astrocytes is increased close to the nucleus and slightly decreased toward the plasma membrane (Figure 5 D). Similarly, we observed a change in the distribution of BCAN in primary cortical *Pclo^gt/gt^* astrocytes (Figure 5 C). In *Pclo^wt/wt^* cortical astrocytes, BCAN-positive puncta are evenly distributed throughout the cytoplasm from the nucleus toward the cell periphery (Figure 5 A). In contrast, BCAN intensity is greater from the nucleus toward the plasma membrane in *Pclo^gt/gt^* astrocytes (Figure 5 A, C), indicating that gene trap Piccolo mutation leads to an altered distribution of BCAN and TNR in cortical astrocytes. Notably, we also observed that BCAN and the TNR were differentially distributed in primary cerebellar astrocytes (Figure S2). Here, BCAN and TNR intensities were slightly greater in *Pclo^gt/gt^* astrocytes along the secretory pathway than in *Pclo^wt/wt^* astrocytes (Figure S2).

**Fig. 5.**
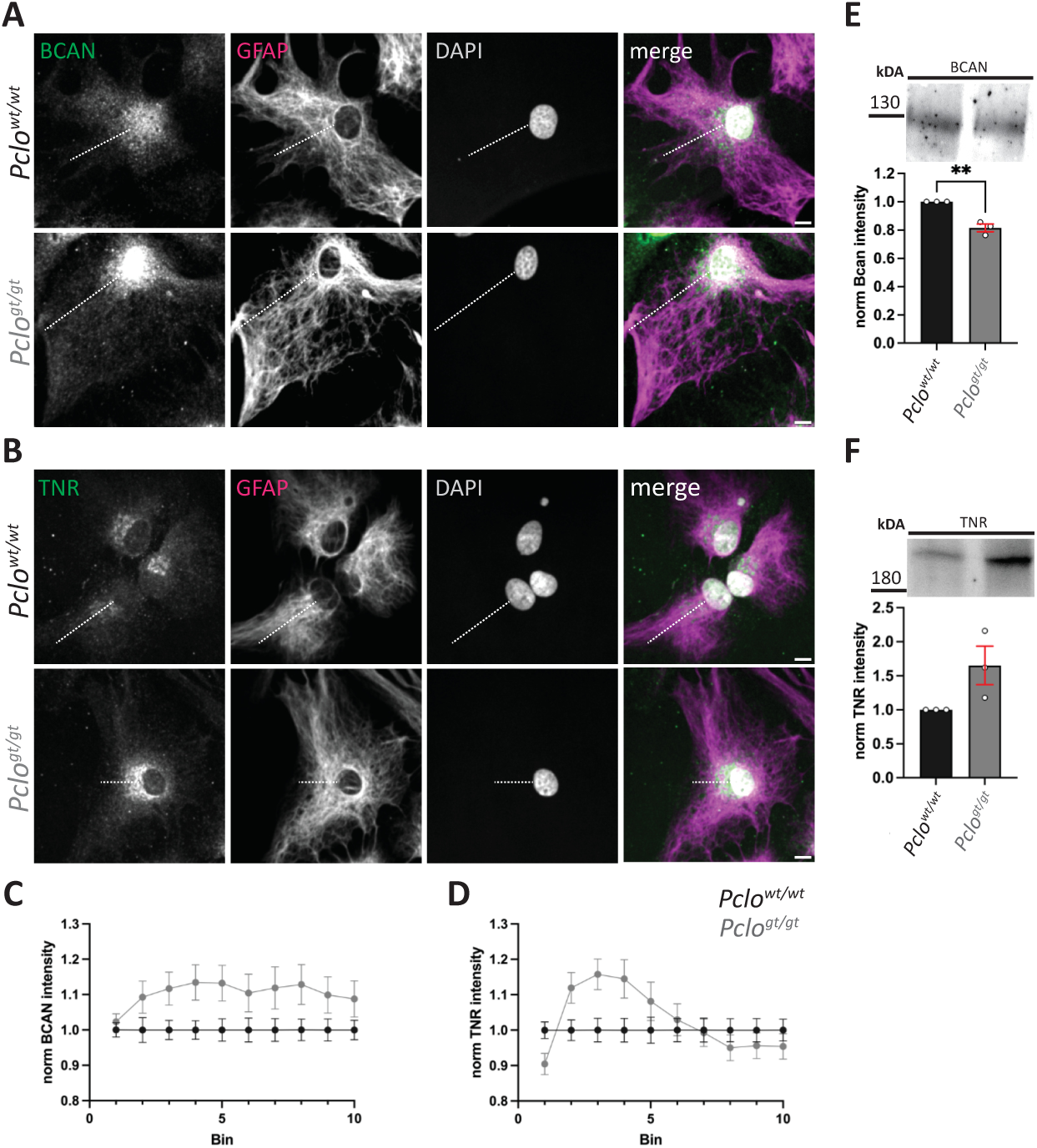
The subcellular distributions of BCAN and TNR are altered in *Pclo ^gt/gt^* primary astrocytes. **A,** Immunocytochemical staining for BCAN on primary *Pclo^wt/wt^* and *Pclo^gt/gt^* cortical astrocytes. In *Pclo^gt/gt^*astrocytes, BCAN intensity accumulates around the nucleus and extends along a line from the center toward the periphery. In *Pclo^wt/wt^* astrocytes, BCAN intensity was distributed more evenly. **B**, Immunocytochemical staining for TNR on *Pclo^wt/wt^*and *Pclo^gt/gt^* cortical astrocytes. In *Pclo^gt/gt^*astrocytes, TNR intensity accumulates around the nucleus and extends along a line from the cell center toward the cell periphery. In contrast, in *Pclo^wt/wt^* astrocytes, the TNR intensity was distributed more evenly. **C**, Quantification of A. A line was drawn from the nucleus to the periphery and divided into 10 bins. BCAN intensity was subsequently measured within each bin and graphically represented. BCAN intensity is greater close to the nucleus in *Pclo^gt/gt^* primary cortical astrocytes than in *Pclo^wt/wt^* primary cortical astrocytes. **D**, Quantification of B. A line was drawn from the nucleus to the periphery and divided into 10 bins, subsequently TNR intensity was measured within each bin. TNR intensity is greater close to the nucleus in *Pclo^gt/gt^* cortical astrocytes than in *Pclo^wt/wt^*. **E**, Western blot (WB) analysis of the supernatants of *Pclo^wt/wt^* and *Pclo^gt/gt^* cortical astrocyte cultures. Significantly less BCAN was detected in *Pclo^gt/gt^* cortical supernatants (*Pclo^wt/wt^*: mean= 1; *Pclo^gt/gt^*: mean± SEM= 0.81± 0.045, n=3 independent experiments; p=0.0024, t test). **F**, Western blot (WB) analysis of the supernatants of *Pclo^wt/wt^*and *Pclo^gt/gt^* cortical astrocyte cultures. Significantly more TNR was detected in supernatant from *Pclo^gt/gt^* cortical astrocytes (*Pclo^wt/wt^*: mean=1, n=3 independent experiments; *Pclo^gt/gt^*: mean± SEM= 1.653± 0.283, n=3 independent experiments; p=0.082, t test). Scale bar, 10 μm.

In addition, we observed that the levels of secreted BCAN and TNR were altered in the supernatants of primary cortical *Pclo^gt/gt^* astrocytes (Figure 5 E, F). Interestingly, the secreted levels of BCAN and TNR exhibit a contrasting pattern, where BCAN levels are reduced (Figure 5 E, *Pclo^wt/wt^*: mean= 1; *Pclo^gt/gt^*: mean± SEM= 0.81± 0.045, n=3 independent experiments; p=0.0024, t test), whereas secreted TNR levels are increased (Figure 5 G, *Pclo^wt/wt^*: mean= 1; *Pclo^gt/gt^*: mean± SEM= 1.653± 0.283, n=3 independent experiments; p=0.082, t test). This phenotype is not restricted to cortical astrocytes but also applies to cerebellar astrocytes (Figure S2). One reason for the altered amounts of secreted BCAN and TNR could be that the numbers of *Pclo^wt/wt^* and *Pclo^gt/gt^* astrocytes are different in culture, for example, due to different growth rates. However, our growth curves revealed that this is not the case (Figure S1), supporting the hypothesis that a shift in Piccolo isoforms leads to alterations in Bcan and TNR secretion from astrocytes.

BCAN can support the formation of synapses and is therefore a main component of PNNs and the perisynaptic ECM, which are known to mediate synaptic plasticity (Nakajo et al. 2025; Dityatev, Frischknecht, and Seidenbecher 2006). Intriguingly, we observed significantly less BCAN at VGlut1 puncta in *Pclo^wt/wt^* neural networks grown together with *Pclo^gt/gt^* than in *Pclo^wt/wt^* astrocytes in a Banker arrangement (Figure 6 A and B, black *^wt^*neuron_*^wt^*astro: mean± SEM= 1± 0.05, n=3 independent experiments; gray *^wt^*neuron_*^gt^*astro: mean± SEM= 0.74± 0.03, n=3 independent experiments; p=0.0001, t test), indicating that impaired astrocytic secretion has a subsequent and direct impact on the formation/composition of perisynaptic nets. In summary, these data support the hypothesis that Piccolo malfunction impairs the secretion/transport of BCAN and TNR from cortical and cerebellar astrocytes. This culminates in a reduced presence of BCAN at perisynaptic nets.

**Fig. 6.**
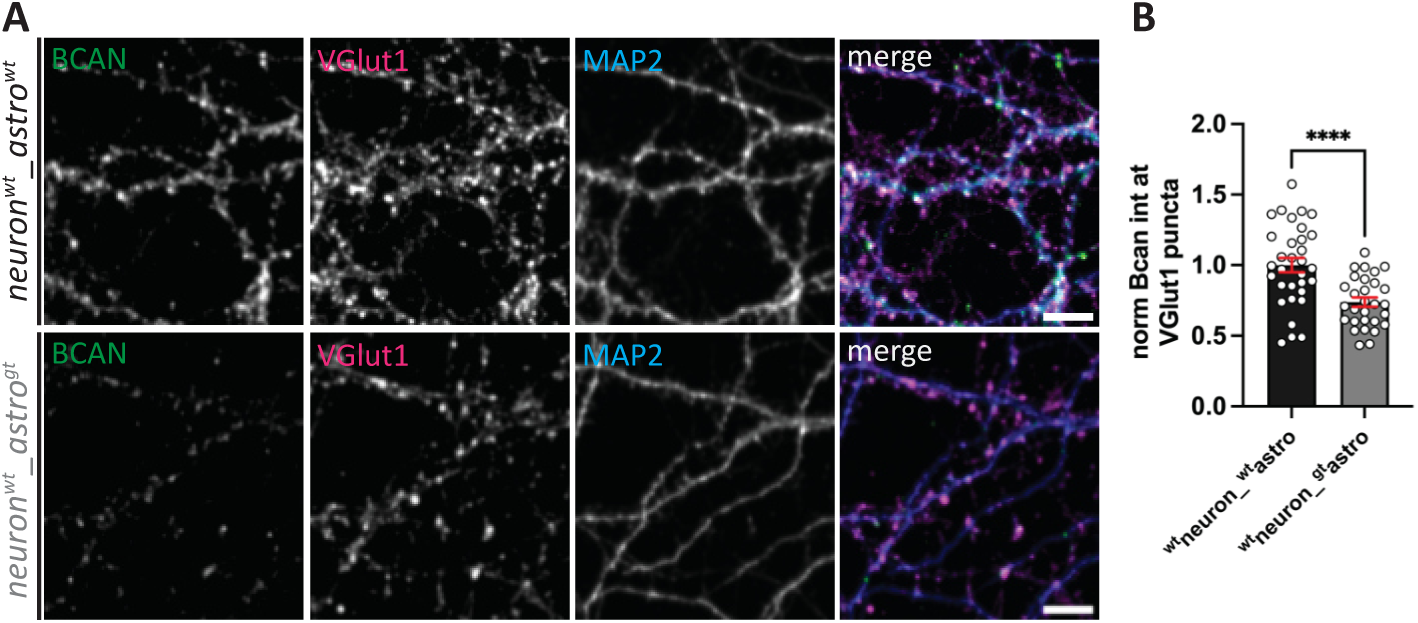
Presence of BCAN at the perisynaptic ECM in neuron networks. **A,** Immunocytochemical staining for BCAN in *Pclo^wt/wt^* or *Pclo^gt/gt^* primary cortical neuron cultures grown together with *Pclo^wt/wt^* or *Pclo^gt/gt^*primary cortical astrocytes. BCAN intensity was subsequently measured at presynaptic terminals marked by VGlut 1. Significantly less BCAN is present at presynaptic terminals in *Pclo^wt/wt^* neuron cultures grown together with *Pclo^gt/gt^* astrocytes. **B,** Quantification of A. BCAN intensity is significantly lower at VGlut1 puncta in *Pclo^wt/wt^*cortical neuron cultures grown together with *Pclo^gt/gt^* cortical astrocytes than in those grown with *Pclo^wt/wt^* cortical astrocytes (black: ^wt^neuron_^wt^astro: mean± SEM= 1± 0.05; gray: ^wt^neuron_^gt^astro: mean± SEM= 0.73± 0.04, n=3 independent experiments; p≤0.0001, t test). Scale bar, 10 μm.

### The morphology of the Golgi apparatus is altered in *Pclo^gt/gt^* primary astrocytes

Protein/cargo transport through the cell is mediated through membrane-enclosed compartments. Secretory cargos move from the ER through the Golgi apparatus to the plasma membrane, where they are released into the extracellular space via membrane fusion (Lippincott-Schwartz, Roberts, and Hirschberg 2000). Here, the Golgi apparatus plays a crucial role within the secretory pathway, as it serves as a hub where secretory cargo proteins are sorted and packaged into specialized “secretory vesicles” for delivery to their specific destinations (Donaldson and Lippincott-Schwartz 2000; Boncompain and Weigel 2018). Owing to the aforementioned changes in the secretion of ECM proteins (Figure 6) and our finding that Piccolo is present at the Golgi in astrocytes (Figure 4), we next analyzed the morphology of the Golgi apparatus in *Pclo^wt/wt^* and *Pclo^gt/gt^* primary astrocytes (Figure 7). Using an antibody against the Golgi protein GM130, we stained primary astrocytes and subsequently analyzed their Golgi structure (Figure 7 A-E). The Golgi structure is more fragmented in *Pclo^gt/gt^* primary cortical astrocytes (Figure 7 A) with numerous small GM130-positive regions (Bins = 1 μm²) (Figure 7 B, C; *Pclo^wt/wt^*: bin1 8.35 bin2 15.52; *Pclo^gt/gt^*: bin1 11.29, bin2 16.93), suggesting that the Golgi integrity is partially impaired upon Piccolo genetrap mutation. In line with this observation, the mean surface fraction area is significantly smaller in *Pclo^gt/gt^*primary cortical astrocytes (Figure 7 A, E; *Pclo^wt/wt^*: median (IQR)= 0.293 (0.675); *Pclo^gt/gt^*: median (IQR)= 0.249 (0.398), 4=independent experiments; p≤0.0001, Mann‒Whitney test). However, the total Golgi area was not significantly different (Figure 7 A and D, *Pclo^wt/wt^*: median (IQR)= 0.942 (0.593); *Pclo^gt/gt^*: median (IQR)= 0.779 (0.636), 4=independent experiments; p=0.052, Mann‒Whitney test). In addition to cortical astrocytes, we observed similar changes in the Golgi apparatus in cerebellar astrocytes (Figure S3). Interestingly, a known interacting partner of Piccolo, PRA1 (S. D. Fenster et al. 2000), localizes to the Golgi in nonneuronal cells and mediates secretion (John et al. 2006). Therefore, we next analyzed immunocytochemical staining for PRA1 and GM130 levels at the Golgi of *Pclo^wt/wt^* and *Pclo^gt/gt^* astrocytes (Figure 7 F and G). We used GM130 staining as a mask and quantified the integrated density (IntDen) of PRA1 per Golgi region (Figure 7 G). Intriguingly, we observed significantly less PRA1 within the Golgi of *Pclo^gt/gt^* astrocytes than in that of *Pclo^wt/wt^* astrocytes. (*Pclo^wt/wt^*: median (IQR)= 0.886 (0.393); *Pclo^gt/gt^*: median (IQR)= 0.711 (0.415), n=3 independent experiments; p=0.0018, Mann‒Whitney Test), indicating that, owing to transposon mutagenesis and a potential change in Piccolo isoforms, less PRA1 is recruited to the Golgi apparatus. PRA1 is known to mediate RabGTPase activity at the golgi (31), interestingly we see altered gene expression of some RAB genes for instance Rab13, Rab32, Rab37, Rab3c, Rab7b, that are present at the golgi (Table 1).

**Fig. 7.**
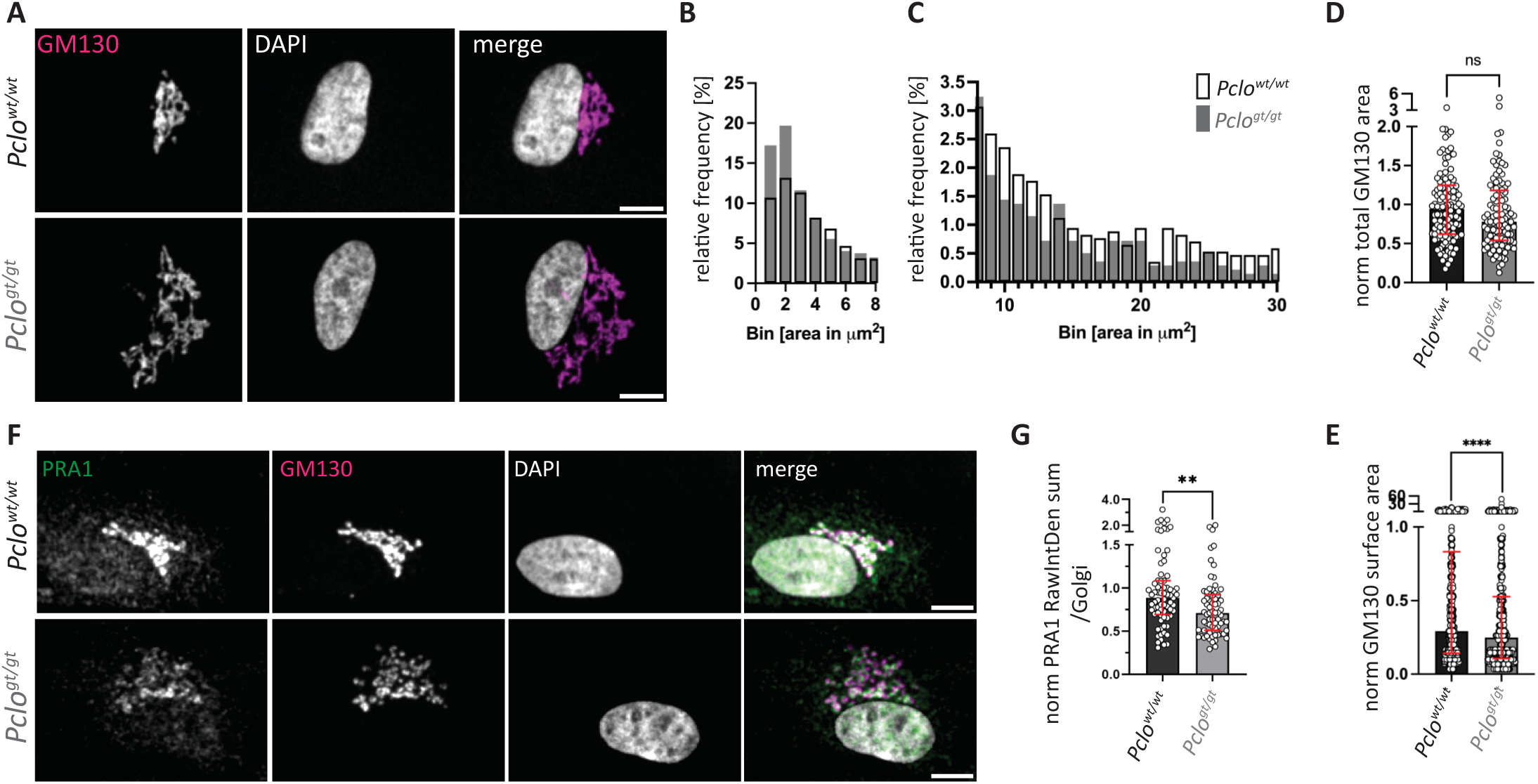
Golgi morphology is altered in *Pclo^gt/gt^* primary cortical astrocytes. **A,** Immunocytochemical staining for GM130 was performed on *Pclo^wt/wt^* and *Pclo^gt/gt^*primary cortical astrocytes from *Pclo^wt/wt^* and *Pclo^gt/gt^*rats. The Golgi morphology slightly differed between *Pclo^wt/wt^*and *Pclo^gt/gt^* astrocytes. Compared to *Pclo^wt/wt^*primary cortical astrocytes, the GM130 staining pattern in *Pclo^gt/gt^*astrocytes appeared more fragmented, with smaller and more dispered areas. **B and C,** Quantification of A. GM130-positive areas were measured, binned by size, and their relative frequency was calculated. In *Pclo^gt/gt^* primary cortical astrocytes, the relative frequency of smaller bins was greater than that in *Pclo^wt/wt^* primary cortical astrocytes. In contrast, the relative frequency of larger fractions was greater in *Pclo^wt/wt^*astrocytes. **D and E**, Quantification of A. The total GM130 area was not significantly different between the *Pclo^wt/wt^* and *Pclo^gt/gt^*Golgi structures (*Pclo^wt/wt^*: median (IQR)= 0.942 (0.593); *Pclo^gt/gt^*: median (IQR)= 0.779 (0.636), n=4 independent experiments; p=0.052, Mann‒Whitney test). However, single GM130 fractions were significantly smaller in *Pclo^gt/gt^* Golgi structures (*Pclo^wt/wt^*: median (IQR)= 0.293 (0.675); *Pclo^gt/gt^*: median (IQR)= 0.249 (0.398), n=4 independent experiments; p≤0.0001, Mann‒Whitney test). Scale bar, 10 μm. **F,** Immunocytochemical staining for PRA1 and GM130 in *Pclo^wt/wt^* and *Pclo^gt/gt^*primary cortical astrocytes**. G,** Quantification of the data in F. GM130 staining was used as a mask and PRA1 integrated density (IntDen) per Golgi area was measured. PRA1 IntDen is significantly lower in *Pclo^gt/gt^* astrocytes than in *Pclo^wt/wt^* astrocytes. (*Pclo^wt/wt^*: median (IQR)= 0.886 (0.393); *Pclo^gt/gt^*: median (IQR)= 0.711 (0.415), n=3 independent experiments; p=0.0018, Mann‒ Whitney test). Scale bar, 10 μm.

**Table 1:** DEGs encoding RabGTPases present in the Golgi apparatus. Pclo gene trap mutation leads to altered gene expression of genes encoding RabGTPases that are linked with Golgi function.

| gene | log2FD change | function |
| --- | --- | --- |
| Rab13 | 0.506 | "late Golgi to post Golgi transport" |
| Rab32 | 0.648 | "post Golgi transport" |
| Rab37 | -1.585 | "secretory autophagy" |
| Rab3c | -0.552 | "KD-Golgi fragmentation" |
| Rab7b | 0.511 | "post Golgi transport" |

Taken together, our data show that Piccolo loss of function leads to altered Golgi morphology and reduced levels of PRA1 in *Pclo^gt/gt^* astrocytes. To date, the role of Piccolo has been described mainly in presynaptic terminals (Wagh et al. 2015; Terry-Lorenzo et al. 2016; Leal-Ortiz et al. 2008; Waites et al. 2013; Ackermann et al. 2019). However, in neurons, Piccolo functions at the Golgi by facilitating the formation and release of Golgi-derived precursor vesicles (PTVs; Piccolo- and Bassoon-positive transport vesicles)(Dresbach et al. 2006; Shapira et al. 2003). Our current results support the hypothesis that Piccolo has an important, more comprehensive role beyond synapses, especially in astrocytes, where it mediates secretion.

### Conditioned media from *Pclo^gt/gt^* astrocytes lead to structural changes in forming neural networks

Astrocytes play a crucial role in regulating brain development and function, as they are, among others, the source of synaptogenic factors (Baldwin and Eroglu 2017) and components of the ECM (Yamada et al. 1997), of which all together mediate synaptogenesis (Baldwin and Eroglu 2017). Therefore, we subsequently investigated the formation of synapses in neural networks (Figure 8). To this end, wild-type primary cortical neurons (*Pclo^wt/wt^*) were cultured with primary cortical astrocytes (*Pclo^wt/wt^* or *Pclo^gt/gt^*) in a Banker setup (e.g., astrocytes on the bottom of plastic Petri dishes and neurons on overhanging glass coverslips), and synapse density (colocalization between synaptophysin and PSD95) was assessed in forming networks (Figure 8 A, B). Notably, synapse density in neural networks grown together with *Pclo^gt/gt^* astrocytes was significantly lower than that in networks grown together with *Pclo^wt/wt^*astrocytes (Figure 8 B, black *^wt^*neuron_*^wt^*astro: mean± SEM= 1± 0.05, n=8 independent experiments; gray *^wt^*neuron_*^gt^*astro: mean± SEM= 0.68± 0.04, n=8 independent experiments; p=0.0001, t test). Similarly, synapse density was also reduced in neural networks grown together with cerebellar *Pclo^gt/gt^* astrocytes (Figure S4 black *^wt^*neuron_*^wt^*astro: mean± SEM= 1± 0.057, n=4 independent experiments; gray *^wt^*neuron_*^gt^*astro: mean± SEM= 0.72± 0.059, n=4 independent experiments; p=0.0001, t test). In addition to immunostaining techniques, we assessed synapse density via electrophysiological recordings. Here, we measured sucrose responses, which reflect the readily releasable pool of vesicles (RRP), in neural mass cultures as a measure of total synapse number (Figure 8 C). We did not observe a significant difference between the RRP sizes of neurons cultured with *Pclo^gt/gt^*astrocytes and those cultured with *Pclo^wt/wt^* astrocytes (Figure 8 C, black astro^wt^_neuron^wt^: median (IQR)= 0.892 (0.656); gray astro^gt^_neuron^wt^: median (IQR)= 0.851 (0.718); n=8 independent experiments, p=0.75; Mann‒Whitney test). Similarly, the RRP of inhibitory neurons cultured with *Pclo^gt/gt^* astrocytes was not significantly different from that of their counterparts grown with *Pclo^wt/wt^* astrocytes (Figure 8 C, black astro^wt^_neuron^wt^: median (IQR)= 0.899 (0.724); gray astro^gt^_neuron^wt^: median (IQR)= 0.870 (0.882); n=8 independent experiments, p=0.68, Mann‒ Whitney test), indicating that the RRP was not significantly affected by the presence of *Pclo^gt/gt^* astrocytes. Taken together, these data suggest that Piccolo malfunction in astrocytes leads to decreased synapse numbers in the formation of neural networks.

**Fig. 8.**
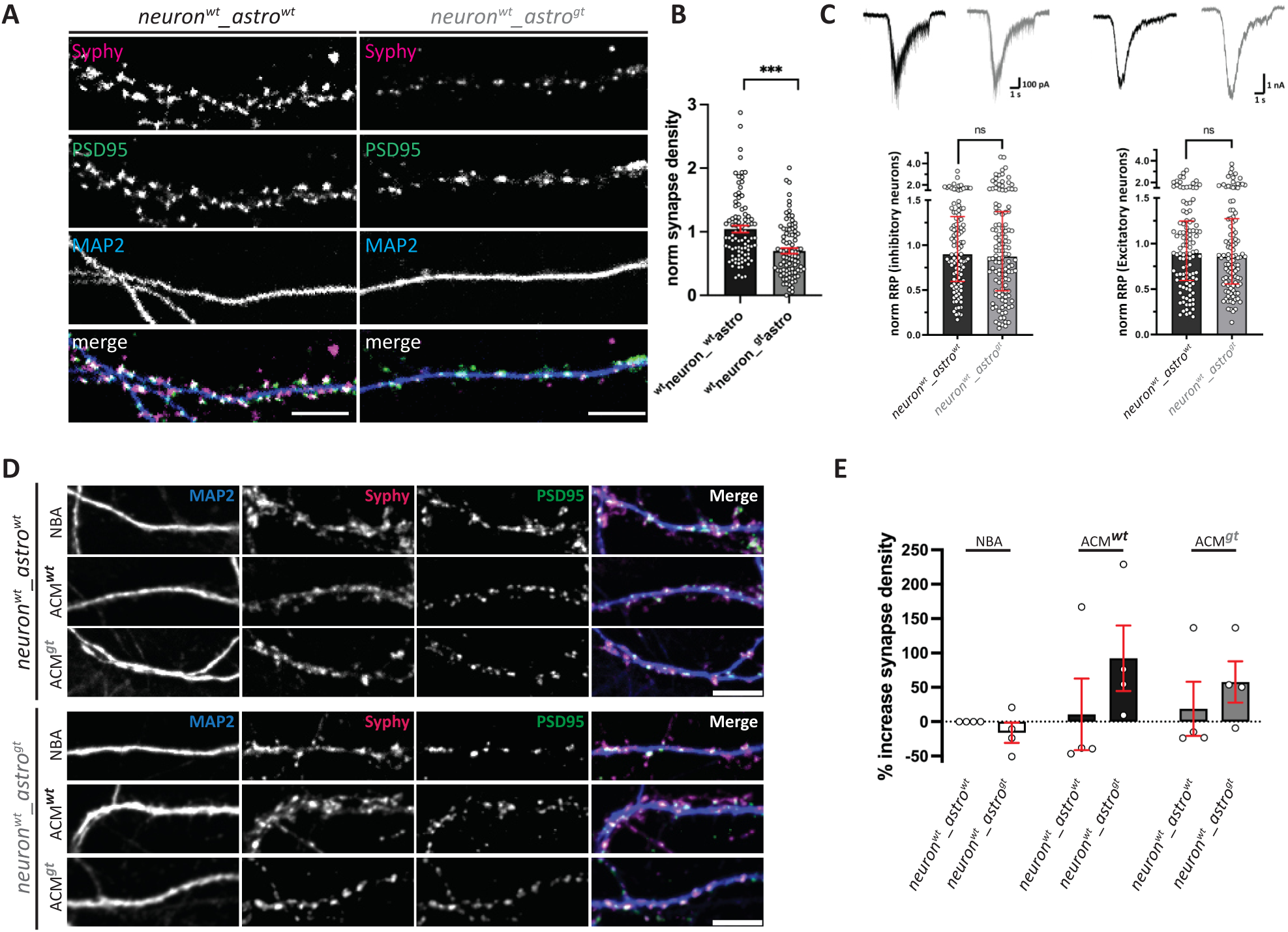
Synapse density is decreased in *Pclo^wt/wt^* cultures grown with *Pclo^gt/gt^*cortical astrocytes. **A**, *Pclo^wt/wt^* or *Pclo^gt/gt^*cortical neuron cultures were grown together with cortical astrocytes (*Pclo^wt/wt^,Pclo^gt/gt^*). Synapse density was assessed by quantifying Syphy-PSD95 double-positive puncta along dendrites (MAP2). In the presence of *Pclo^gt/gt,^*astrocyte synapse density is reduced. **B**, Quantification of A. Synapse density is significantly reduced in the presence of *Pclo^gt/gt^*astrocytes (black: ^wt^neuron_^wt^astro: mean± SEM= 1± 0.051; gray: ^wt^neuron_^gt^astro: mean± SEM= 0.68± 0.044, n=8 independent experiments; p≤0.0001, t test). **C,** Measurement of RRPs. Top: representative traces of RRP response to 0.5 M sucrose in excitatory (left) and inhibitory (right) neurons cultured with *Pclo^wt/wt^*(black) or *Pclo^gt/gt^* (gray) astrocyte. Bottom left: quantification of RRP responses from excitatory neurons (*Pclo^wt/wt^*: median (IQR)= 0.892 (0.656); *Pclo^gt/gt^*: median (IQR)= 0.851 (0.718); n=8 independent experiments, p=0.75, Mann‒Whitney test). Bottom right: quantification of RRP responses from inhibitory neurons (*Pclo^wt/wt^*: median (IQR)= 0.899 (0.724); *Pclo^gt/gt^*: median (IQR)= 0.870 (0.882); n=8 independent experiments, p=0.68; Mann‒Whitney test). **D,** *Pclo^wt/wt^* cortical neuron cultures were grown together with *Pclo^wt/wt^* or *Pclo^gt/gt^* astrocytes and supplemented with NBA, ACM^wt^ or ACM^gt^. Synapse density is reduced in *Pclo^wt/wt^* neuron – *Pclo^gt/gt^* astrocyte cocultures supplemented with NBA. It increased in the presence of ACM^wt^. Only a small increase in synapse density was observed in the presence of ACM^gt^. **E**, Quantification of d. Synapse density is reduced in *Pclo^wt/wt^* neuron – *Pclo^gt/gt^* astrocyte cocultures supplemented with NBA (astro^wt^_neuron^wt^: 0; astro^gt^_neuron^wt^: - 16.11±14.82, n=4 independent experiments). An increase was observed in the presence of ACM^wt^ (astro^wt^_neuron^wt^: mean± SEM= 34.11± 49.87, astro^gt^_neuron^wt^: mean± SEM= 92.19± 47.67; n=4 independent experiments). A small increase was observed in the presence of ACM^gt^ (astro^wt^_neuron^wt^: mean± SEM= 18.65± 39.38, astro^gt^_neuron^wt^: mean± SEM= 38.63± 16.08; n=4 independent experiments). Scale bar, 10 μm.

### Astrocyte-conditioned media restores synapse density in neural networks cultured with *Pclo^gt/gt^* astrocytes

Importantly, our neural networks were established in a “Banker setup” (Banker and Goslin 1988), wherein astrocytes were grown on the bottom of a plastic dish and neurons were grown on a suspended glass cover slip. As such, the two cell types are not in physical contact with each other. This arrangement supports the hypothesis that the phenotypes we observed in neural networks juxtaposed with *Pclo^gt/gt^*astrocytes are attributable to defects in astrocytic secretion. This finding is further supported by the fact that the addition of ACM^wt/wt^ (<u>a</u>strocyte-<u>c</u>onditioned <u>m</u>edia) significantly increased synapse density in neural networks grown together with *Pclo^gt/gt^* astrocytes. ACM^wt/wt^ addition increased synapse density in neural networks grown together with *Pclo^gt/gt^*astrocytes by 90 % compared to neural networks grown with *Pclo^wt/wt^* astrocytes in pure NBA media. In contrast, it only induced the formation of 34% additional synapses in neural cultures grown together with *Pclo^wt/wt^* astrocytes (Figure 8 D, E ACM^wt/wt^ astro^gt^_neuron^wt^: mean± SEM= 92.19± 47.67; n=4 independent experiments; ACM^wt/wt^ astro^wt^_neuron^wt^: mean± SEM = 34.11± 49.87, n=4 independent experiments). Notably, the addition of ACM^gt/gt^ also led to increased synapse density in neural networks grown together with *Pclo^gt/gt^* astrocytes; however, this density was lower than that in cultures receiving ACM^wt/wt^. An increase in synapse density of only ∼20% was observed in ACM^gt/gt^-treated neural networks grown together with *Pclo^wt/wt^* astrocytes (Figure 8 D, E ACM^gt/gt^ astro^gt^_neuron^wt^: mean± SEM= 38.63± 16.08, n=4 independent experiments; ACM^gt/gt^ astro^wt^_neuron^wt^: mean± SEM= 18.65± 39.38, n=4 independent experiments).

Taken together, our ACM^wt/wt^ and ACM^gt/gt^ data support the hypothesis that *Pclo^gt/gt^* astrocytes secrete reduced levels of synapse-promoting factors. Intriguingly, we see gene expression changes of other genes coding for synaptogenic factors like Bdnf, Semaphorin, Thrombospondin and Ephrin (Table 2).

**Table 2:**
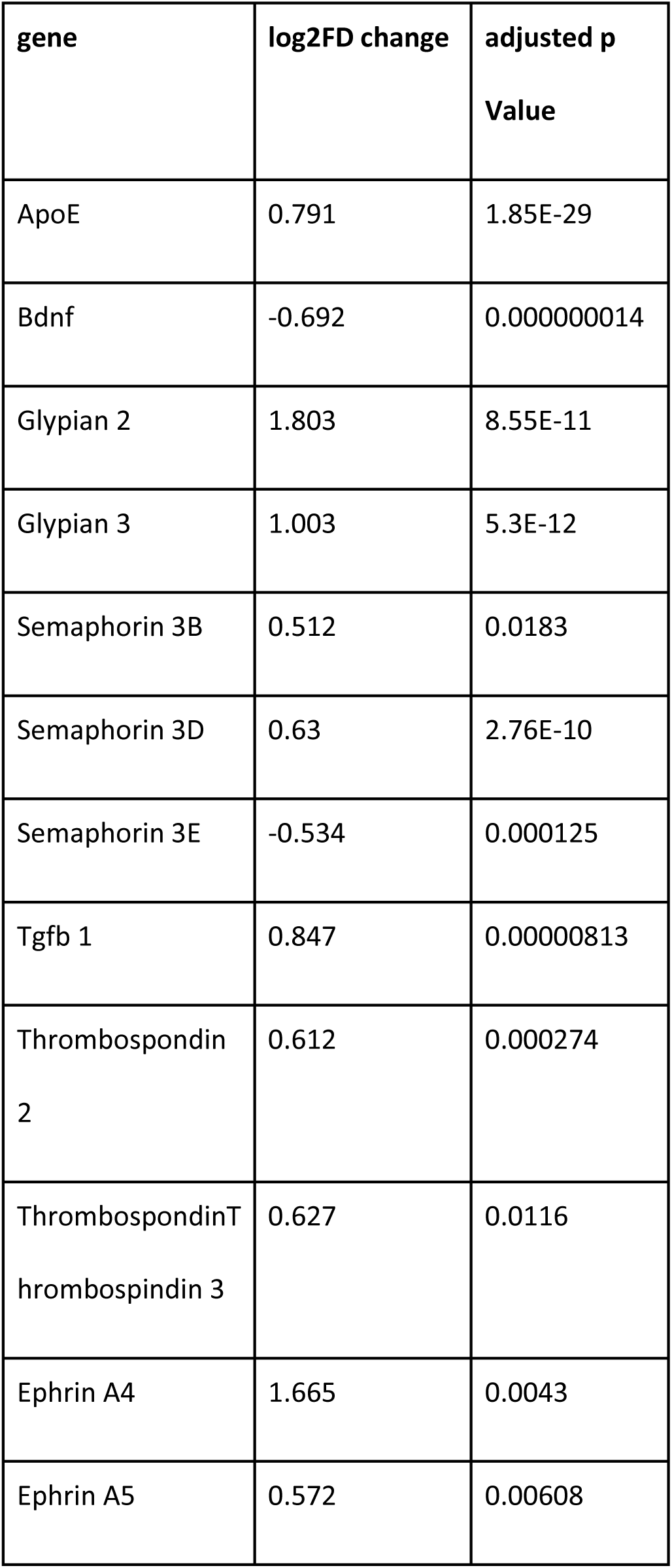
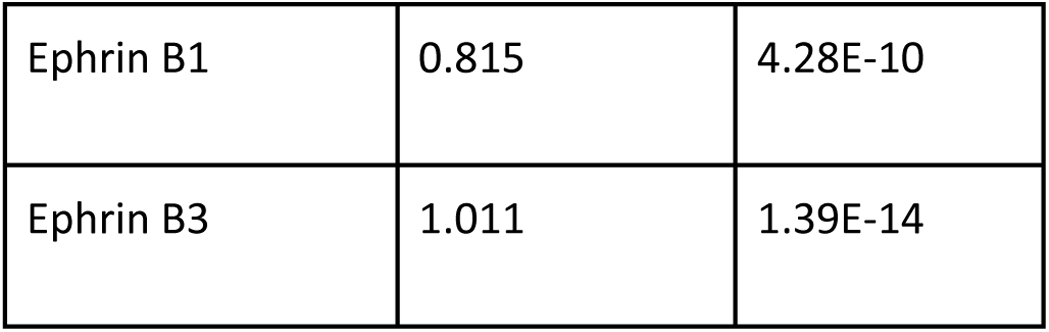
DEGs encoding synaptogenic factors. Pclo gene trap mutation leads to altered gene expression of genes coding for synaptogenic factors that become secreted from astrocytes.

### *Pclo^gt/gt^* astrocytes lead to functional changes in neural networks

Alterations in synapse density within neural networks, as we observed in *Pclo^wt/wt^* networks grown together with *Pclo^gt/gt^*astrocytes (Figure 8), can have a tremendous effect on network function, especially their activity and neural output. To test this concept, we next analyzed spontaneous network activity by recording miniature postsynaptic currents and compared neural cultures grown with *Pclo^wt/wt^*astrocytes with those grown with *Pclo^gt/gt^* astrocytes (Figure 9). We distinguished between excitatory and inhibitory cell types by measuring both miniature excitatory postsynaptic currents (mEPSCs) and miniature inhibitory postsynaptic currents (mIPSCs) in our cultures. Compared with those grown with *Pclo^wt/wt^* astrocytes, wild-type neuron mass cultures grown together with *Pclo^gt/gt^* astrocytes presented a significant increase in mEPSC frequency (Figure 9 C, black astro^wt^_neuron^wt^: median (IQR)= 0.826 (0.934); gray astro^gt^_neuron^wt^: median (IQR)= 1.046 (1.324); n=8 independent experiments, p=0.021; Mann‒Whitney test). In addition, mEPSC amplitudes were not significantly different in cultures grown with *Pclo^gt/gt^* astrocytes than in those grown with *Pclo^wt/wt^* astrocytes (Figure 9 D black astro^wt^_neuron^wt^: median (IQR)= 0.937 (0.452); gray astro^gt^_neuron^wt^: median (IQR)= 1.007 (0.632); n=8 independent experiments, p=0.082; Mann‒Whitney test). In contrast to the mEPSC frequency, we observed no difference in the mIPSC frequency in neuron cultures grown together with *Pclo^gt/gt^* compared with those grown with *Pclo^wt/wt^* astrocytes (Figure 9 black astro^wt^_neuron^wt^: median (IQR)= 0.808 (1.019); gray astro^gt^_neuron^wt^: median (IQR)= 0.715 (0.85); p=0.518, n=8 independent experiments; Mann‒Whitney test). Notably, the mIPSC amplitude of neuron cultures grown together with *Pclo^gt/gt^* or *Pclo^wt/wt^*astrocytes was not significantly different (Figure 9 black astro^wt^_neuron^wt^: median (IQR)= 0.938 (0.535), n=122 data points; gray astro^gt^_neuron^wt^: median (IQR)= 891 (0.652); n=8 independent experiments; p=0.863, Mann‒Whitney test). Thus, our recordings indicate that the presence of *Pclo^gt/gt^* astrocytes can lead to an increase in the intrinsic excitability of neural networks.

**Figure 9:**
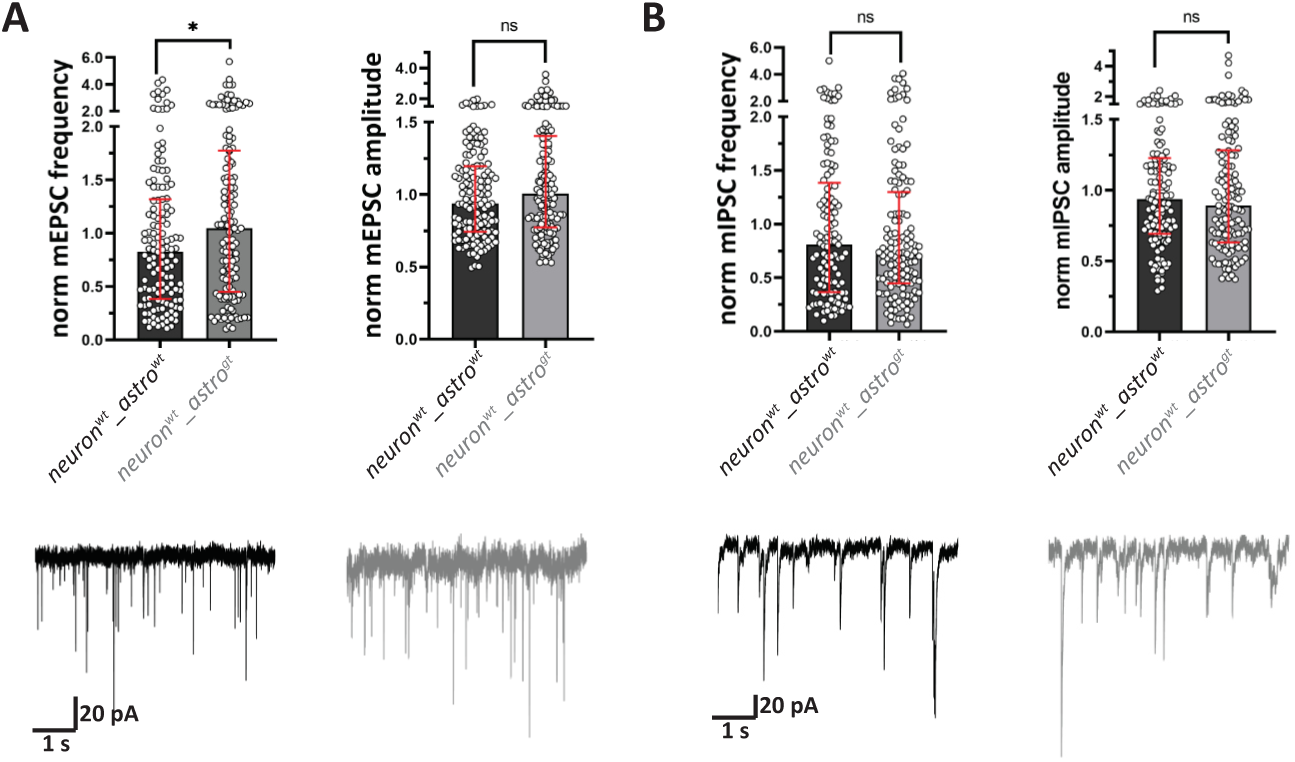
Neuronal excitability is altered in *Pclo^gt/gt^* cultures grown together with *Pclo^gt/gt^* cortical astrocytes. **A,** Miniature excitatory postsynaptic currents (mEPSCs) were measured in neurons cultured in Banker setups with either *Pclo^wt/wt^* (black) or *Pclo^gt/gt^*(gray) astrocytes. The mEPSC frequency significantly increased in cultures supported by *Pclo^gt/gt^* astrocytes (astro^wt^_neuron^wt^: median (IQR)= 0.826 (0.934); astro^gt^_neuron^wt^: median (IQR)= 1.046 (1.324); n=8 independent experiments, p=0.021; Mann‒Whitney test). In contrast, there was no significant difference in mEPSC amplitude between the two culture conditions. (astro^wt^_neuron^wt^: median (IQR)= 0.937 (0.452); astro^gt^_neuron^wt^: median (IQR)= 1.007 (0.632); n=8 independent experiments, p=0.082, Mann‒ Whitney Test). **B,** Miniature inhibitory postsynaptic currents (mIPSCs) were also measured from neuron cultures grown together with either *Pclo^wt/wt^*or *Pclo^gt/gt^* astrocytes. There was no significant difference in mIPSC frequency in cultures supported by *Pclo^wt/wt^* astrocytes or *Pclo^gt/gt^* astrocytes (astro^wt^_neuron^wt^: median (IQR)= 0.808 (1.019); astro^gt^_neuron^wt^: median (IQR)= 0.715 (0.85); n=8 independent experiments, p=0.518; Mann‒Whitney test). Similarly, the mIPSC amplitude did not differ significantly between the cultures (astro^wt^_neuron^wt^: median (IQR)= 0.938 (0.535); astro^gt^_neuron^wt^: median (IQR)= 891 (0.652); n=8 independent experiments, p=0.863; Mann‒Whitney test). Scale bar, 10 μm.

Taken together, our data demonstrate that *Pclo^gt/gt^* astrocytes influence the activity of excitatory neurons, which will have a direct effect on the excitation‒inhibition balance and therefore the activity and output of neural networks.

## Discussion

In this study, we revealed a crucial role for the scaffolding protein Piccolo in astrocytes, which regulates BCAN and TNR secretion to shape synapse formation. While Piccolo has previously been associated with various presynaptic functions, its role in astrocytes has remained unidentified until now.

In presynaptic terminals, Piccolo’s functions are linked primarily to membrane trafficking pathways such as synaptic vesicle recycling and protein degradation, in addition to regulating actin dynamics (Wagh et al. 2015; Leal-Ortiz et al. 2008; Waites et al. 2013; Ackermann et al. 2019). Our findings demonstrate that Piccolo’s function in astrocytes is also closely tied to membrane dynamics and secretion. Looking at two secretory cargos, BCAN and TNR, we show that Piccolo regulates secretion from astrocytes, which impacts downstream formation of neuronal networks (Figures 7, 8, and 9). This novel discovery positions Piccolo as a regulator of membrane trafficking across various cell types, including neurons and astrocytes, with significant implications for synapse and neural circuit formation. Notably, Piccolo plays a crucial yet underexplored role in the development of neurodegenerative diseases such as PCH3. Our findings revealed that changes in Piccolo isoform distribution, impaired synapse formation and activity—an early pathological event among others that may trigger disease progression. Given its pivotal role, future research should prioritize uncovering the precise mechanisms by which Piccolo contributes to neurodegenerative disease pathogenesis, potentially opening new avenues for therapeutic intervention.

### The phenotypes resulting from Piccolo loss of function become increasingly pronounced as postnatal development progresses

Earlier studies analyzing *Pclo^gt/gt^* rats demonstrated that Piccolo loss of function results in a complex, multilayered phenotype that impacts various brain regions and cell types (Rajab et al. 2003; Ahmed et al. 2015; Falck et al. 2020). Interestingly, our current study revealed a significant increase in the number of DEGs at later developmental stages in both cerebellar and brainstem tissues (Figure 1). These findings suggest that the effects of Piccolo loss of function worsen over time as the animals reach the juvenile stage. Notably, children with PCH3 are born without apparent abnormalities but begin to exhibit typical phenotypes at the age of 5 to 1 months (Ahmed et al. 2015). Since P25 is still an early stage in a rat’s lifespan, future studies should investigate how the RNA-seq profile evolves in old, respectively aging rats to better understand later developmental stages and their implications for PCH3 manifestation.

### Piccolo loss of function results in defects in the secretion of ECM components from astrocytes

Given the relatively small number of DEGs identified in young rats (P5), we focused our subsequent analysis on DEGs found in the P25 cerebellum and brainstem tissue. Interestingly, GO enrichment analysis revealed that many of the proteins encoded by these DEGs are linked to key biological processes such as signaling, cell communication, and the ECM (Figure 2). These processes are well-known regulators of neural outgrowth, synaptogenesis, and synapse function (Jiang, Sando, and Südhof 2021; Scheiffele 2003; Yang et al. 2023; Ferrer-Ferrer and Dityatev 2018), and disruptions in these signaling cascades can lead to altered synapse morphology and function, as previously described in *Pclo^gt/gt^* cerebella (Falck et al. 2020). Of particular interest are changes in ECM components (such as BCAN and TNR), bone morphogenetic proteins (BMPs), thrombospondins and semaphorins, which are crucial modulators of synaptogenesis and synapse function (Baldwin and Eroglu 2017). Our data revealed a significant reduction in the distribution of BCAN outside astrocytes in *Pclo^gt/gt^* brain sections as well as in the supernatants from primary astrocyte cultures (Figures 3, 5, S3). BCAN, along with other ECM components such as aggrecan, is a key constituent of the perisynaptic ECM (Faissner et al. 2010; Dityatev, Seidenbecher, and Schachner 2010; Frischknecht and Seidenbecher 2012). The perisynaptic ECM forms directly at presynaptic terminals (Frischknecht and Seidenbecher 2012). Interestingly, in neural networks grown together with *Pclo^gt/gt^* astrocytes, BCAN levels are significantly reduced at VGlut1-positive presynaptic terminals (Figure 6), further supporting the hypothesis that the secretion of BCAN from *Pclo^gt/gt^*astrocytes is impaired. Reduced levels of BCAN may stem from two possible mechanisms: (1) impaired secretion of BCAN from astrocytes and/or (2) increased degradation of already secreted BCAN within the extracellular space. In support of the first hypothesis, we observed disrupted BCAN distribution along the secretory trafficking route from the ER/Golgi toward the cell periphery in *Pclo^gt/gt^* primary cortical and cerebellar astrocytes (Figures 5, S3). This alteration indicates that loss of Piccolo function can negatively affect BCAN intracellular transport, impairing its secretion from astrocytes (Figures 5, S3), which supports the hypothesis that Piccolo is essential for BCAN secretion from cortical and cerebellar astrocytes.

An important related question is whether this effect is specific to BCAN or extends to other secretory ECM cargoes. Interestingly, we found that the TNR distribution is also altered along the secretory pathway in *Pclo^gt/gt^* astrocytes and in their supernatants but in an opposite manner from that of BCAN, e.g., more TNR is detected in *Pclo^gt/gt^*cell culture supernatants than in *Pclo^wt/wt^* cell culture supernatants (Figure 5). These findings suggest that a shift in Piccolo isoforms in astrocytes affects several aspects of the secretory pathway and not just a single cargo pathway. Notably, the distribution of nonsecreted signaling receptors, such as BMPR1b, remains unchanged in *Pclo^gt/gt^*astrocytes (data not shown), suggesting that Piccolo isoforms in *Pclo^gt/gt^* astrocytes disrupt the trafficking of secretory cargoes without affecting nonsecretory cargoes. However, the secreted levels of BCAN and TNR are altered in opposing directions (Figure 5). Cargo secretion follows a common pathway from the ER through the Golgi and trans-Golgi network to the plasma membrane, where it is released into the extracellular space via membrane fusion (Cunliffe 2005). However, different cargoes take distinct trafficking routes depending on their secretion mechanisms. Broadly, secretion is classified into lysosomal (mainly for enzymes), regulated, and constitutive pathways (Kelly 1985; Néel et al. 2024). The different effects on BCAN and TNR secretion in *Pclo^gt/gt^* astrocytes likely arise from their distinct secretion mechanisms. For example, tenascin C (TNC) is secreted by astrocytes in a regulated manner upon serum stimulation (Nishio et al. 2003), suggesting that TNR may follow a similarly regulated secretion pathway. In contrast, BCAN might be secreted constitutively.

While Piccolo’s role in astrocyte secretion is novel and not previously described, it has been linked to insulin secretion from pancreatic beta cells, linking it to this process in other cell types (Fujimoto et al. 2002). In this study, we identified, for the first time, Piccolo isoforms in astrocytes whose domain structure is distinct from that of neurons expressing full-length Piccolo. We looked at the c-terminal domains PDZ and C2A and found changes in their structure (Figure 4). With respect to the C2A domain, we amplified an additional smaller band from mRNAs isolated from astrocytes, which represents a differentially spliced C2A domain that lacks exon 16 and has been described previously in mice (Figure 4, (Garcia et al. 2004)). Interestingly, the long C2A domain, which is found in neurons, displays Ca^2+^-dependent dimerization accompanied by phospholipid binding (Garcia et al. 2004). In contrast, the smaller C2A domain, which we find in astrocytes, does not result in a Ca^2+^-dependent conformational change; however, its Ca^2+^ affinity and therefore phospholipid binding are increased (Garcia et al. 2004). Similarly, we observed different alternatively spliced versions of the PDZ domain in neurons and astrocytes (Figure 4). Here, we observed a double band in neurons, again indicating that alternatively spliced isoforms of Piccolo exist in neurons. Among those, only the smaller band was present in *Pclo^wt/wt^*astrocytes (Figure 4). Generally, a shift in the distribution of Piccolo isoforms between neurons and astrocytes suggests that Piccolo performs different functions in each cell type. The presence of the smaller C2A domain suggests that piccolo may have different activities in astrocytes because of its higher Ca^2+^ affinity (Garcia et al. 2004; Gerber et al. 2001), and alternatively spliced PDZ domains could impact binding to interaction partners and therefore subsequent cellular functions. Intriguingly, gene trap transposon mutagenesis in *Pclo^gt/gt^* rats induces an additional shift in the distribution of astrocytic Piccolo isoforms (Figure 4), which could have an impact on function. In *Pclo^gt/gt^* astrocytes, Piccolo isoforms that harbor both a long and a short C2A domain are still present; however, the ratio of these isoforms is altered (Figure 4), which could influence proper function. The shift in splicing and Piccolo isoform distribution is particularly pronounced for the PDZ domain. Here, only the small version exists in *Pclo^wt/wt^* astrocytes, whereas the larger as well as the smaller isoform is present in *Pclo^gt/gt^* astrocytes (Figure 4). A shift in isoform distribution can have a direct effect on potential interaction partners of Piccolo in *Pclo^wt/wt^* and *Pclo^gt/gt^* astrocytes and therefore on its cellular function. Here, the cAMP GFF Epac2 is of specific interest; it forms with Piccolo via its PDZ domain a functional complex in pancreatic islet cells, which mediates Ca2+ dependent insulin secretion (Fujimoto et al. 2002).

The role of Piccolo in astrocytes has not been described previously. Its subcellular localization can indicate its function; Piccolo predominantly localizes to the Golgi apparatus in astrocytes but is also detected in a punctate pattern in the juxtaposed cytoplasm (Figure 4). Both Piccolo and its close homolog bassoon harbor a Golgi-binding domain (GBR), through which they localize to the Golgi in neurons (Dresbach et al. 2006; Tao-Cheng 2007). Thus, Piccolo isoforms could also bind to the Golgi via this domain in astrocytes (Figure 4). In neurons, Piccolo and Bassoon mediate the formation of active zone precursor transport vesicles (PTVs) from the Golgi to nascent synapses (Maas et al. 2012). It is therefore feasible that Piccolo could mediate the formation of a subclass of secretory vesicles from the Golgi in astrocytes. In particular, the trans Golgi (TNG) contains adapter proteins such as AP1, which help to select and recruit cargo molecules into secretory vesicles (Tan and Gleeson 2019). As a scaffolding protein that binds phospholipids in a Ca^2+^-dependent manner (Gerber et al. 2001), Piccolo could select cargo molecules, similar to AP1, by interacting with distinct cargo molecules, for instance, via its PDZ domain. Changes in the PDZ domain structure could cause a change in binding affinity to interaction partners of Piccolo at the Golgi in *Pclo^gt/gt^* astrocytes. This could influence cargo selection as well as the formation of functional protein complexes and thereby the secretion of BCAN and TNR (Figure 3, 5). In parallel, the observed shift in the expression of Piccolo isoforms harboring long or short C2A domains could affect Ca^2+^-dependent signaling and phospholipid binding of Piccolo in *Pclo^gt/gt^* astrocytes. Notably, many processes within the Golgi, such as secretory vesicle budding, are Ca^2+^-dependent (Sargeant and Hay 2022), and a shift in the Ca^2+^ affinity of Piccolo’s C2A domain could influence these processes. Furthermore, malfunction in membrane flux, the Golgi transport machinery, or the cytoskeletal interaction system, have been linked with Golgi morphology (Zappa, Failli, and De Matteis 2018). Indeed, we observed an abnormal Golgi morphology in *Pclo^gt/gt^* astrocytes, where the cisternae appeared less compact and more fragmented (Figure 6, Figure S4). Piccolo is known to interact with components of the cytoskeleton (Terry-Lorenzo et al. 2016) in neurons similarly it could link it with the Golgi in astrocytes. The observed change in Piccolo isoforms in *Pclo^gt/gt^* astrocytes could interfere with this function and cause Golgi fragmentation. Notably, the known Piccolo interaction partner PRA1 also localizes to the Golgi in astrocytes; however, its levels are lower in *Pclo^gt/gt^* astrocytes than in *Pclo^wt/wt^* astrocytes (Figure 7). Notably, PRA1 interacts with Rab5 and other Rab GTPases, promoting their activation. At presynaptic boutons, Piccolo tethers PRA1 near the endocytic machinery, mediating Rab5 activation and endosomal membrane trafficking (Ackermann et al. 2019). Piccolo could also mediate Rab GTPase activity via PRA1 at the Golgi and therefore play a role in secretory vesicle trafficking. As less PRA1 is recruited to the Golgi in *Pclo^gt/gt^* astrocytes, this could have an impact on secretory cargo trafficking. Moreover, our RNA-seq data revealed altered expression of multiple Rab GTPase-encoding genes (Rab13, Rab32, Rab37, Rab3c, and Rab7b) (Table 1). Interestingly, Rab3c-knockdown cells exhibit severe Golgi fragmentation (Galea and Simpson 2015), mirroring the phenotypes observed in *Pclo^gt/gt^* astrocytes (Figure 6, S4). Notably, Golgi fragmentation is an early phenotype observed in neurodegenerative diseases such as Alzheimer’s disease and Parkinson’s disease (Martínez-Menárguez et al. 2019). Thus, future experiments are necessary to further explore the relationships among Piccolo, Golgi function and secretion from astrocytes.

### The loss of Piccolo function in astrocytes disrupts the formation and function of synapses

ECM components, including BCAN and TNR, are well-established synaptogenic factors that regulate synapse formation and function (Frischknecht and Seidenbecher 2012; Dityatev and Schachner 2003; Bukalo, Schachner, and Dityatev 2007). They are, along with other ECM components such as aggrecan, key constituents of perisynaptic nets (Faissner et al. 2010; Frischknecht and Seidenbecher 2012; Chelini et al. 2024). Unlike perineuronal nets (PNNs), which mainly enclose inhibitory interneurons, the perisynaptic ECM forms directly at presynaptic terminals (Frischknecht and Seidenbecher 2012). This raises a critical question: do the observed changes in BCAN/TNR secretion from *Pclo^gt/gt^* astrocytes as well as the reduced levels of BCAN at presynaptic terminals affect synaptogenesis and neural circuit development? Indeed, we found evidence of impaired synaptogenesis in our Piccolo gene trap rat model (Falck et al. 2020). In this study, primary cortical *Pclo^wt/wt^* neurons that were grown together with *Pclo^gt/gt^* astrocytes formed significantly fewer synapses than those grown with *Pclo^wt/wt^*. This strongly suggests that, owing to impaired secretion of for instance BCAN and/or TNR from *Pclo^gt/gt^* astrocytes, synaptogenesis is decreased. This finding is compounded by data showing that *Pclo^wt/wt^*astrocyte-conditioned media can rescue the phenotype (Figure 8). We cannot, however, conclude that altered levels of BCAN or TNR are exhaustively responsible. However, BCAN-KO mice show changes in signal transmission as well as a significant reduction in VGlut1 puncta at the Calyx of Held (Blosa et al. 2015), indicating that the observed loss of BCAN at perisynaptic sites can lead to the formation of fewer synapses. We cannot rule out that other synaptogenic factors, which are also secreted from astrocytes and whose gene expression is altered in *Pclo^gt/gt^*brain tissue, contribute to the observed synapse density phenotype.

In addition to structural changes, an altered synaptic ECM composition can also impact neural network activity. Intriguingly, BCAN-KO mice show changes in fast signal transmission as well as the maintenance of long-term potentiation (Blosa et al. 2015; Brakebusch et al. 2002) in the hippocampus. Similarly, we observed a significant increase in mEPSC frequency in neural networks growing together with *Pclo^gt/gt^*astrocytes compared with those growing with *Pclo^wt/wt^* astrocytes, whereas the mIPSC frequency did not change (Figure 9). These findings suggest that networks grown together with *Pclo^gt/gt^*astrocytes have greater intrinsic excitability, which could be due to synaptic changes as well as overall network activity. This could affect neural output, contributing to the phenotypes observed in PCH3 patients and *Pclo^gt/gt^*rats; it could particularly explain seizures that are present in PCH3 patients and *Pclo^gt/gt^* rats (Ahmed et al. 2015; A. Medrano et al. 2018).

As previously mentioned, BCAN and TNR are most likely not the only signaling factors whose secretion is affected in *Pclo^gt/gt^* astrocytes. In addition to ECM components, astrocytes secrete additional synaptogenic factors that coordinate and support synapse formation (Baldwin and Eroglu 2017). For example, thrombospondin, EphrinA3 and BDNF are critical regulators of excitatory synapse formation (Baldwin and Eroglu 2017). Interestingly, the gene expression of these synaptogenic factors is also altered in *Pclo^gt/gt^* cerebellum tissue (Table 2), which could explain why *Pclo^gt/gt^* rats present defects in synapse formation and maturation. Future experiments will be necessary to determine the exact secretome of *Pclo^gt/gt^*astrocytes and thus affect signaling pathways, opening new avenues for the development of potential treatments.

## Conclusion

This study reveals a crucial and previously unrecognized role for the scaffolding protein Piccolo in astrocytes, where it mediates the secretion of the ECM components BCAN and TNR that are essential for neuronal network formation. We demonstrate that Piccolo mutation, as modeled in our *Pclo^gt/gt^* rat, leads to impaired ECM secretion, resulting in reduced synapse density and dysregulated activity in neuronal networks. These findings establish a novel cellular and molecular mechanism linking astrocytic Piccolo dysfunction to the pathology of Pontocerebellar Hypoplasia Type 3 (PCH3) and other neurodegenerative conditions. Our work further highlights the critical role of astrocytes in neuronal circuit integrity and suggests new avenues for astrocyte-targeted therapeutic strategies.

## List of abbreviations

AALAC: Assessment and Accreditation of Laboratory Animal Care International
AD: Alzheimer Disease
ACM: astrocyte-conditioned media
AMBIO: Advanced Medical Bioimaging Core Facility
BCAN: brevican
BDNF: brain derived neurotrophic factor
BMPR1b: bone morphogenetic protein receptor type 1B
BP: biological process
CC: cellular component
CHMP4C: charged multivesicular body protein 4c
DEG: differentially expressed gene
ECM: extracellular matrix
ER: endoplasmatisches Retikulum
FDR: false discovery rate
GBR: golgi-binding domain
GC: granula cell
GCL: granula cell layer
GFAP: glial fibrillary acidic protein
GM130: golgi-matrix-protein 130 kDa
GO: gene ontology
IACUC: Institutional Animal Care Use Committee
IQR: interquartile Range
mEPSC: mini excitatory postsynaptic current
MF: mossy fiber
mIPSC: mini inhibitory postsynaptic current
MOF: molecular function
Pclo: Piccolo
PCH3: Pontocerebellar Hypoplasia Type 3
PD: Parkinson’s Disease
PRA1: pernylated Rab acceptor 1
PSD95: postsynaptic density protein 95
PTV: Piccolo-and Bassoon-positive transport vesicles
SLC25A25: solute carrier family 25 member 25
SNP: single-nucleotide polymorphism
SV: synaptic vesicle
TBC1D4: TBC1 domain family member 4
TNG: trans golgi network
TNR: tenascin-R
TNC: tenascin C
TNG: trans golgi network
RRP: readily releasable pool
TTX: tetrodotoxin

## Declarations

### Ethics approval

All procedures for experiments involving animals, were approved by the animal welfare committee of Charité Medical University and the Berlin state government.

### Consent for publication

not applicable.

### Availability of the data and material

The datasets used and/or analysed during the current study are available from the corresponding author on reasonable request.

## Acknowledgements

We thank the Advanced Medical Bioimaging Core Facility (AMBIO) of the Charité – Universitätsmedizin for support in the acquisition of imaging data.

---

**S1.**
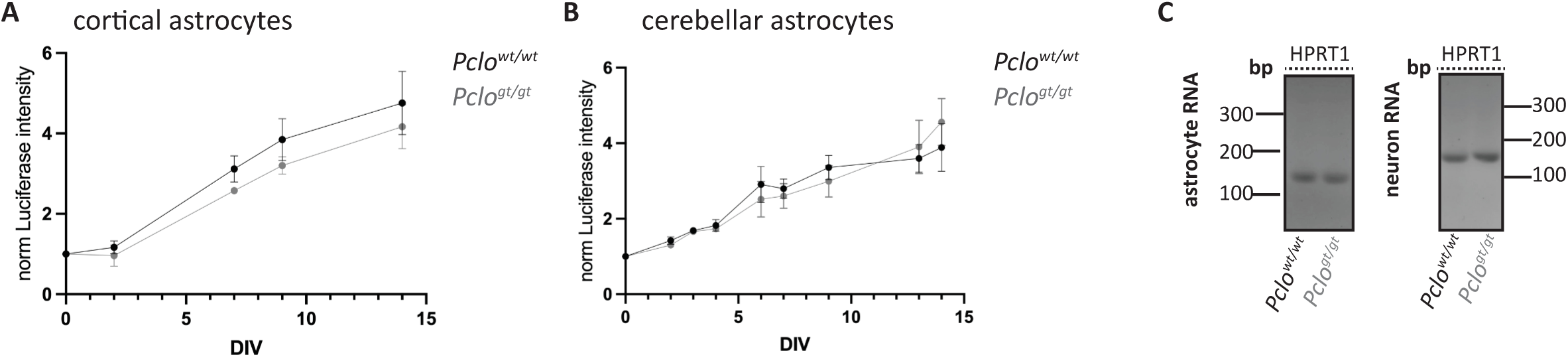
*Pclo^wt/wt^*and *Pclo^gt/gt^* astrocytes show similar growth rates. **A-B**, Astrocyte growth rates were determined by measuring luciferase activity every other day for 15 days. *Pclo^wt/wt^* and *Pclo^gt/gt^* astrocytes grew at the same growth rate, and no significant difference was detected. **C,** RT‒PCR on mRNA isolated from *Pclo^wt/wt^* and *Pclo^gt/gt^*cortical astrocytes as well as cortical neurons. Primers for the housekeeping gene *HPRT1* were used to assess RNA integrity..A clear HPRT1 band was amplified from all samples, confirming good RNA quality.

**S2:**
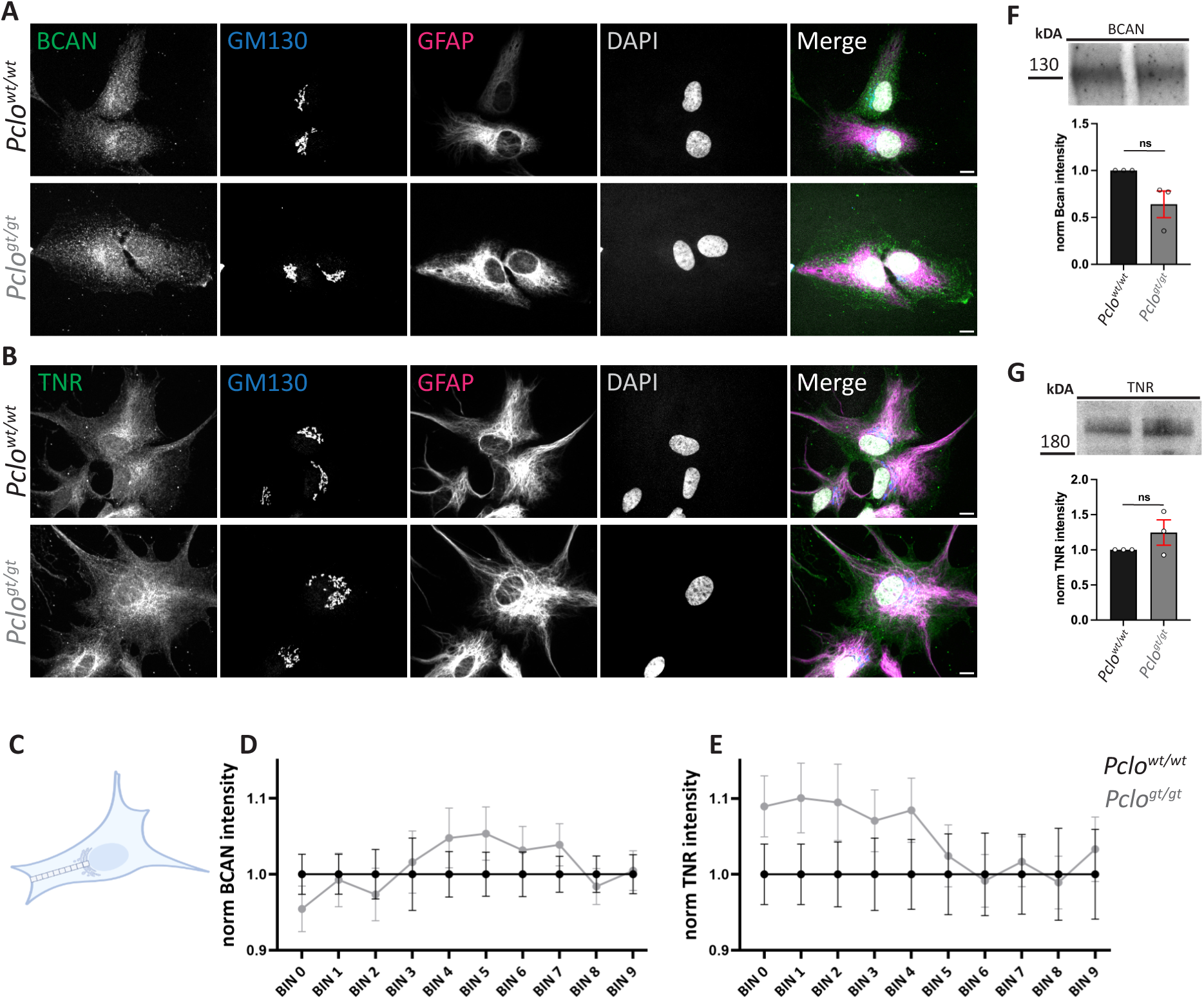
Subcellular distribution of BCAN and TNR in *Pclo^wt/wt^* and *Pclo ^gt/gt^*cerebellar astrocytes. **A,** Immunocytochemical staining BCAN, GM130, GFAP, and DAPI in *Pclo^wt/wt^* and *Pclo ^gt/gt^* cerebellar astrocytes. BCAN fluorescence intensity is diffusely distributed in *Pclo^wt/wt^*astrocytes, whereas it accumulates along a defined path in *Pclo ^gt/gt^* astrocytes**. B,** Immunocytochemical staining of tenascin-R (TNR), GM130, GFAP, and DAPI in *Pclo^wt/wt^* and *Pclo ^gt/gt^* cerebellar astrocytes. TNR fluorescence intensity is concentrated near the nucleus and extends toward the cell periphery in *Pclo ^gt/gt^* astrocytes, in contrast to a more uniform distribution in *Pclo^wt/wt^* astrocytes. **C,** Schematic illustration of the cargo distribution analysis method. A line from the nucleus to the cell periphery was segmented into 10 equal bins (0–9), and the fluorescence intensity of the cargo was measured in each bin and normalized to the mean intensity per bin in *Pclo^wt/wt^* astrocytes. **D,** Quantification of a. Normalized BCAN intensity decreases progressively from the nucleus to the periphery in *Pclo^wt/wt^* astrocytes but slightly increases in *Pclo ^gt/gt^* astrocytes. **E,** Quantification of B. Normalized TNR intensity is greater near the nucleus in *Pclo ^gt/gt^* astrocytes than in *Pclo^wt/wt^*cells. **F,** Western blot (WB) analysis of BCAN levels in the supernatants of *Pclo^wt/wt^* and *Pclo^gt/gt^* cerebellar astrocytes. BCAN levels are reduced in *Pclo^gt/gt^* astrocytes (*Pclo^wt/wt^*: mean= 1; *Pclo^gt/gt^*: mean± SEM= 0.641± 0.141, n=3 independent experiments; p=0.064, t test). **G,** Western blot (WB) analysis of TNR levels in the supernatants of *Pclo^wt/wt^* and *Pclo ^gt/gt^*cerebellar astrocytes. TNR levels are increased in *Pclo^gt/gt^* astrocytes (*Pclo^wt/wt^*: mean= 1; *Pclo^gt/gt^*: mean± SEM= 1.246± 0.179, n=3 independent experiments; p=0.242, t test). Scale bar, 10 μm.

**S3.**
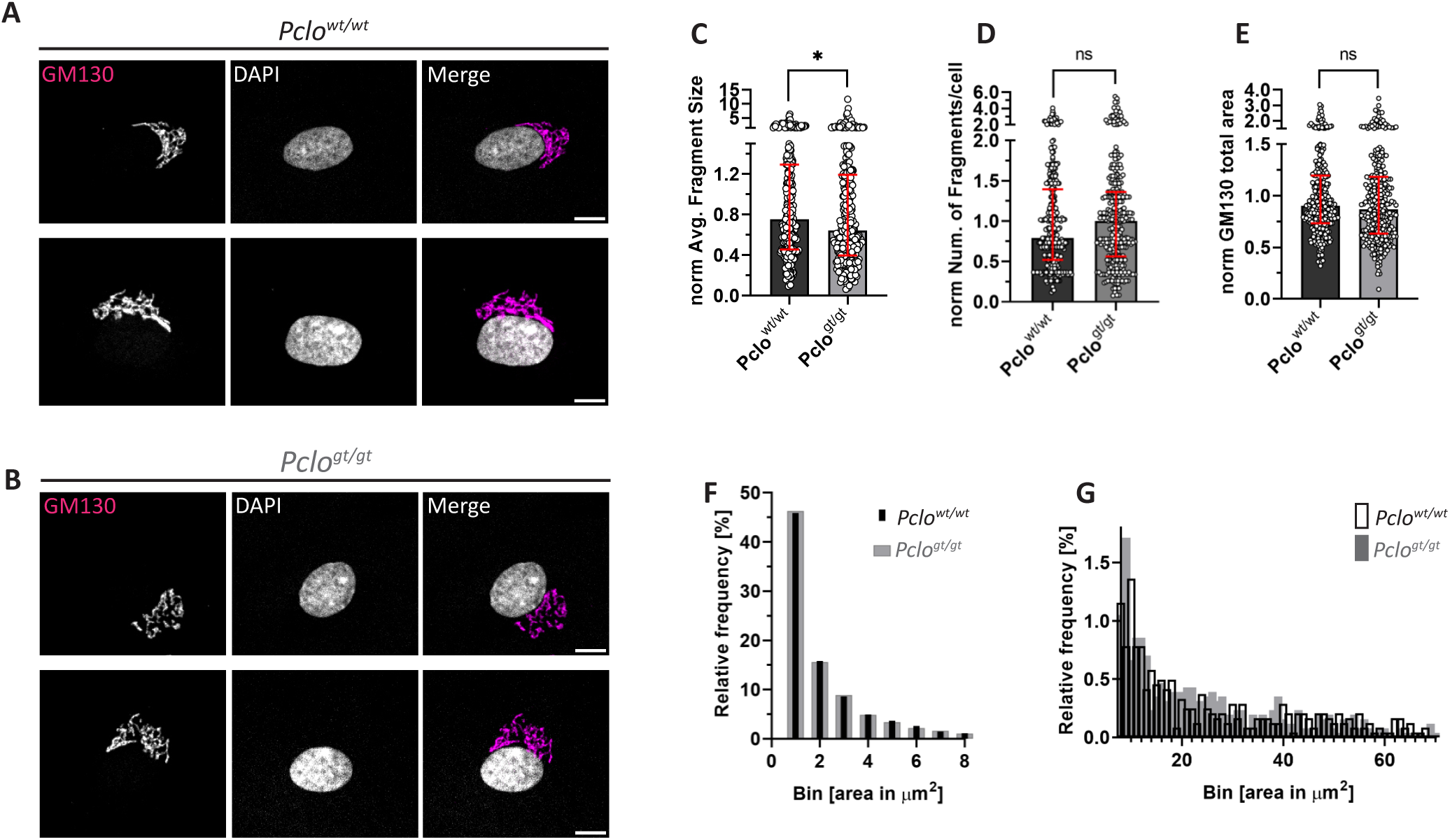
Golgi morphology is altered in *Pclo^gt/gt^* cerebellar astrocytes. **A and B,** Representative immunocytochemical staining of GM130 in primary cerebellar astrocytes from *Pclo^wt/wt^* (A) and *Pclo^gt/gt^* (B) rats. Note the apparent fragmentation of the Golgi apparatus in *Pclo^gt/gt^*astrocytes, which are characterized by smaller and more dispersed GM130-positive areas than *Pclo^wt/wt^* astrocytes. **C-E,** Quantitative analysis of GM130 staining. **C**, Average size of GM130-positive fragments per cell, normalized to the mean of *Pclo^wt/wt^*, showing a significant decrease in *Pclo^gt/gt^* astrocytes (*Pclo^wt/wt^*median (IQR)= 0.75 (0.84), n=307 data points; *Pclo^gt/gt^* median (IQR)= 0.640 (0.79), n=310 data points; p=0.0199). **D**, Number of GM130-positive fragments per cell, normalized to the mean of *Pclo^wt/wt^*, showing a trend toward an increase in *Pclo^gt/gt^* astrocytes (*Pclo^wt/wt^* median (IQR)= 0.79 (0.87), *Pclo^gt/gt^* median (IQR)= 1 (0.8),p=0.099). **E,** Total GM130-positive area per cell, normalized to the mean of *Pclo^wt/wt^*, showing a slight decrease in *Pclo^gt/gt^*astrocytes (*Pclo^gt/gt^* median (IQR)= 0.90 (0.46), *Pclo^gt/gt^*median (IQR)= 0.87 (0.55); p=0.113. **F, G** Relative frequency distribution of GM130-positive fragment sizes. **F,** Frequency of small fragments (0–8 µm²). **G,** Frequency of large fragments (≥9 µm²). All the data were collected from n=4 independent experiments, and statistical significance was determined via the Mann‒Whitney U test, with p< 0.05 considered significant. Scale bar, 10 μm.

**S4.**
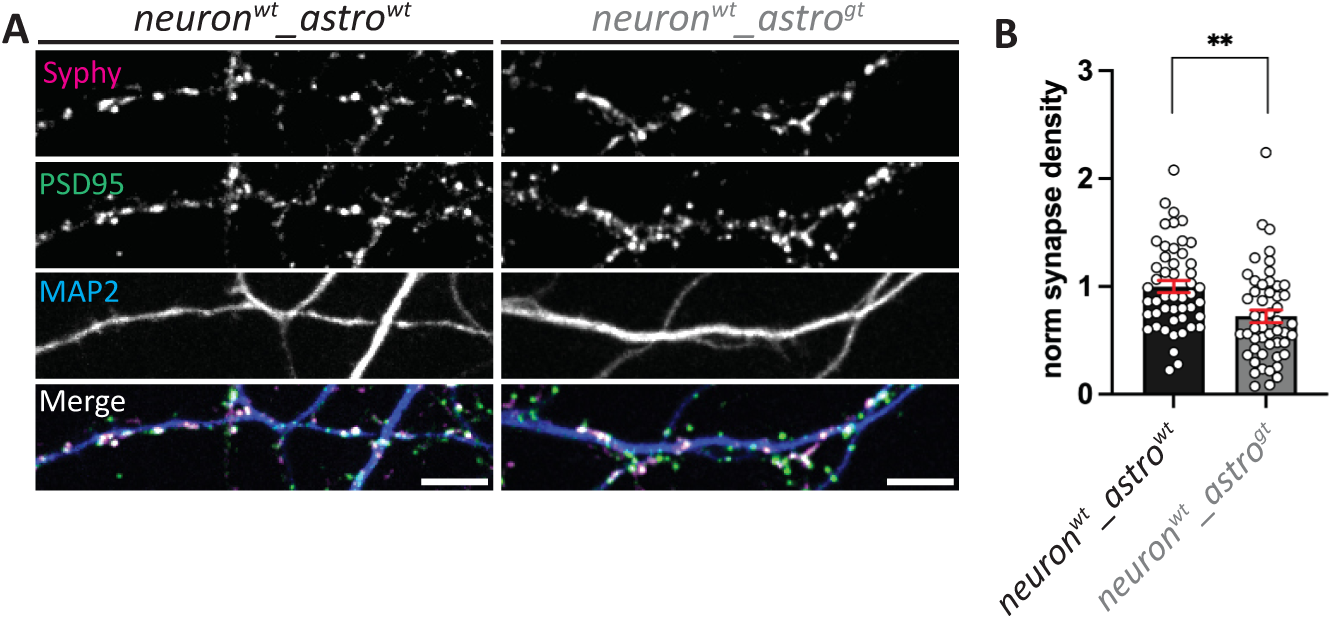
Synapse density is decreased in *Pclo^wt/wt^* cultures grown together with *Pclo^gt/gt^*cerebellar astrocytes. **A**, *Pclo^wt/wt^* or *Pclo^gt/gt^* primary cortical neuron cultures were grown together with *Pclo^wt/wt^* or *Pclo^gt/gt^* primary cerebellar astrocytes. Synaptic density was subsequently measured by counting Syphy - PSD95 double-positive puncta along MAP2-positive dendrites. In the presence of *Pclo^gt/gt,^* cerebellar astrocyte synapse density was reduced. **B**, Quantification of A. Synapse density was significantly lower in *Pclo^wt/wt^* cortical neuron cultures grown together with *Pclo^gt/gt^* cerebellar astrocytes than in those grown with *Pclo^wt/wt^* cerebellar astrocytes (black, gray; ^wt^neuron_^wt^astro mean± SEM= 1± 0.057; ^wt^neuron_^gt^astro mean± SEM= 0.72± 0.059, n=4 independent experiments; p≤0.0011, t test). Scale bar, 10 μm.

